# Individual variation in the functional lateralization of human ventral temporal cortex: Local competition and long-range coupling

**DOI:** 10.1101/2024.10.15.618268

**Authors:** Nicholas M. Blauch, David C. Plaut, Raina Vin, Marlene Behrmann

## Abstract

The ventral temporal cortex (VTC) of the human cerebrum is critically engaged in high-level vision. One intriguing aspect of this region is its functional lateralization, with neural responses to words being stronger in the left hemisphere, and neural responses to faces being stronger in the right hemisphere; such patterns can be summarized with a signed laterality index (LI), positive for leftward laterality. Converging evidence has suggested that word laterality emerges to couple efficiently with left-lateralized frontotemporal language regions, but evidence is more mixed regarding the sources of the right-lateralization for face perception. Here, we use individual differences as a tool to test three theories of VTC organization arising from: 1) local competition between words and faces driven by long-range coupling between words and language processes, 2) local competition between faces and other categories, 3) long-range coupling with VTC and temporal areas exhibiting local competition between language and social processing. First, in an in-house functional MRI experiment, we did not obtain a negative correlation in the LIs of word and face selectivity relative to object responses, but did find a positive correlation when using selectivity relative to a fixation baseline, challenging ideas of local competition between words and faces driving rightward face lateralization. We next examined broader local LI interactions with faces using the large-scale Human Connectome Project (HCP) dataset. Face and tool LIs were significantly anti-correlated, while face and body LIs were positively correlated, consistent with the idea that generic local representational competition and cooperation may shape face lateralization. Last, we assessed the role of long-range coupling in the development of VTC lateralization. Within our in-house experiment, substantial positive correlation was evident between VTC text LI and that of several other nodes of a distributed text-processing circuit. In the HCP data, VTC face LI was both negatively correlated with language LI and positively correlated with social processing in different subregions of the posterior temporal lobe (PSL and STSp, respectively). In summary, we find no evidence of local face-word competition in VTC; instead, more generic local interactions shape multiple lateralities within VTC, including face laterality. Moreover, face laterality is also influenced by long-range coupling with social processing in the posterior temporal lobe, where social processing may become right-lateralized due to local competition with language.

## 1 Introduction

Visual recognition is a fundamental skill performed by humans and animals alike, important both in its own right and for guiding higher-order interactions with inanimate objects and social agents. The neural substrate of visual recognition is implemented in the ventral visual pathway and is comprised of a hierarchy of regions that transform retinal inputs into high-level features suitable for recognition across large naturalistic variation in the inputs (Mishkin et al., 1983; Felleman and Van Essen, 1991; DiCarlo and Cox, 2007; Yamins and DiCarlo, 2016b). After processing in the early retinotopic areas, signals are propagated to the *ventral temporal cortex* (VTC), a large extrastriate area encompassing multiple stages of mid- and high-level visual processing.

Many investigations have characterized the functional organization of VTC—how representations of stimulus and task properties are spatially organized across the cortical surface (Kanwisher, 2010). For example, in almost all participants, a face-preferring region can be found on the fusiform gyrus (Kanwisher et al., 1997; Chen et al., 2023), a word-preferring region can be found near the occipitotemporal sulcus (McCandliss et al., 2003; Rueckl et al., 2015), and a place-preferring region can be found near the parahippocampal gyrus (Epstein et al., 1999) and collateral sulcus (for review, see Grill-Spector and Weiner, 2014). In addition to the presence of category-selective areas, a different perspective has noted super-areal spatial organization for higher-order properties, such as animacy and real-world size (Konkle and Caramazza, 2013), or, more abstractly, dimensions of object space (Bao et al., 2020). Within this object space perspective, the first two dimensions of organization were proposed to be animacy and stubbiness/spikiness (Bao et al., 2020); however, one study, controlling for image-based confounds, has suggested that the distinction between faces and bodies holds greater importance in the organization of human VTC than stubiness/spikiness (Yargholi and Op de Beeck, 2023). Recently, work has demonstrated that both the low dimensions of object space (Bao et al., 2020), as well as the higher dimensions are topographically organized in the macacque inferotemporal cortex (Yao et al., 2023), in line with other research demonstrating hierarchical topographic organization of different properties within VTC (Tanaka, 1996, 2003; Brants et al., 2011; Sato et al., 2013). Many aspects of VTC topographic organization have recently been shown to emerge within self-organizing neural network models, given pressures to perform high-level object recognition tasks under spatial pressures to keep connectivity short or to exhibit local correlation (Blauch et al., 2022b; Doshi and Konkle, 2023; Margalit et al., 2024).

## 1.1 Faces and words in VTC

Humans are expert at recognizing faces, a skill acquired over evolutionary timescales (Parr, 2011). As such, the presence of one or more visually face-selective areas in VTC has been seen as tantalizing support of a possible adaptive and evolved functional network (Kanwisher, 2010; Ratan Murty et al., 2020). However, most humans are also expert at the visual recognition of words, a skill which has only become widespread in most societies over the last few hundred years (Roser and Ortiz-Ospina, 2018), which is too short of a timeline for human brains to have evolved a particular mechanism for this ability (Dehaene and Cohen, 2007; Polk and Farah, 1998). Nevertheless, a word-selective region emerges in VTC in virtually all readers (McCandliss et al., 2003; Dehaene et al., 2010), but is absent in illiterate individuals (Dehaene et al., 2010). That VTC contains a response profile selective for words is strong evidence for the idea that experience-dependent plasticity can drive the large-scale functional organization of VTC to reflect the visual statistics of ecologically relevant tasks (Dehaene and Cohen, 2007; Yamins and DiCarlo, 2016a).

Importantly, faces and words demonstrate complementary lateralization patterns, with face selectivity typically stronger in the right hemisphere (RH), and word selectivity typically stronger in the left hemisphere (LH). Rather than being confined to a single selective area, there are multiple areas that respond preferentially for each of these categories within VTC in many or most observers. For face processing, there is general consensus of the presence of an occipital face area, OFA (Gauthier et al., 2000), and two fusiform face-selective areas: FFA-1 and FFA-2 (also referred to as pFus-faces and mFus-faces) in the posterior and middle fusiform gyrus, respectively (Grill-Spector et al., 2017; Weiner et al., 2017). These regions are generally considered to be part of a *core* hierarchical network (Gobbini and Haxby, 2007), homologous to the macaque face-patch system that has been studied in greater detail (Tsao et al., 2003, 2008; Bao et al., 2020). For words, it is thought that there are posterior regions in each hemisphere, L-VWFA-1 and R-VWFA-1, that support perceptual feature extraction, and a more anterior and left-dominant region, L-VWFA-2, that communicates with downstream language areas (Lerma-Usabiaga et al., 2018; White et al., 2019). White et al. (2019) has suggested that interhemispheric communication prior to VWFA-2 allows for the combination of word-related information from each visual hemifield to converge in the left VWFA-2, where word recognition is accomplished. Additionally, face and word recognition have each been viewed as recruiting an *extended* distributed cortical network, with regions in VTC being seen as critical nodes involved in higher-level feature extraction, part of the general processing demands for each of these categories (Gobbini and Haxby, 2007; Rosenthal et al., 2017; Bouhali et al., 2014; Stevens et al., 2017; Vin et al., 2024).

## 1.2 Individual variability

The functional organization of VTC is broadly consistent across individuals, as reflected in the success of group-level analyses. However, examination of individual functional maps also reveals substantial inter-participant variability. Indeed, this variability has motivated the widespread use of functional localizers that are designed to demarcate a selective cluster of voxels within an anatomical search space so that the response profile of the functionally-defined region can be analyzed on a single-participant basis (Kanwisher et al., 1997; Kanwisher, 2010). This approach circumvents the spatial blurring of heterogeneous functional regions across individuals; in the case of the VWFA, individual variability obscures the observation of word-selective responses in VTC using a purely anatomical definition (Glezer and Riesenhuber, 2013).

Somewhat surprisingly, however, the individual variability in spatial organization that motivates the use of functional localizers—variability not only in the location, but also in the size, shape, and hemispheric organization of functional regions, as well as in the number of functional regions—has received only limited consideration. This has painted a somewhat rigid view of the mosaic of functional topography in VTC, emphasizing a principled developmental outcome rather than a principled developmental process. Notably, although the classic VWFA (VWFA-2) is typically left-lateralized, it exhibits a rightward laterality in participants exhibiting rightward language dominance in frontotemporal areas (Cai et al., 2008, 2010; Van der Haegen et al., 2012; Gerrits et al., 2019). This finding would be expected if VWFA inherits its laterality through functional coupling with downstream language lateralization(Behrmann and Plaut, 2020), and preferential anatomical and functional connectivity between VWFA and language areas (Bouhali et al., 2014; Saygin et al., 2016; Stevens et al., 2017). Moreover, recent work has demonstrated clear individual differences in both the size and number of face-selective regions in VTC (Gao et al., 2022; Chen et al., 2023). Overall, the marked variability challenges the claim of canonical areal organization (Felleman and Van Essen, 1991; Kanwisher, 2010), suggesting a more stochastic developmental process with multiple possible local and global solutions in terms of areal organization and inter-areal connectivity (Sporns et al., 2004; Astle et al., 2023). In this way, individual variability may provide clues into the developmental origins of VTC organization (Davies-Thompson et al., 2016; Dundas et al., 2015).

## 1.3 Competition between words and faces?

What gives rise to the topographic and hemispheric organization of words and faces? The *neuronal recycling* (NR) hypothesis (Dehaene and Cohen, 2007) suggests that the VWFA emerges in a specific LH region that both receives foveal inputs and has connectivity with LH language regions, satisfying the input and output demands of reading. A related theory of *graded hemispheric specialization* (GHS) has proposed that the acquisition of literacy creates a competitive pressure with faces in the LH VTC, which leads to the rightward lateralization of face processing (Plaut and Behrmann, 2011; Behrmann and Plaut, 2015). Specifically, both words and faces compete to be proximal to high-acuity visual information in lateral cortical regions, with words competing more strongly in the LH to also be proximal to left-lateralized language areas. Critical to GHS, this competition is not monolithic and varies across individuals, resulting in graded degrees of lateralization (Behrmann and Plaut, 2020). Objects, which are recognized with less expertise and thus place less demand on foveal resources, and scenes, which place greater demands on peripheral vision, are thus proposed to be largely spared from this competition, in line with their weaker lateralization (Hasson et al. 2002, for related ideas, see Levy et al. 2001).

One important difference between NR and GHS is that, while GHS agrees that neuronal recycling may be at play, it considers cortical competition more generally, encompassing both *recycling* (i.e. one category “stealing” the resouces of another) and *blocking* (i.e. preventing the acquisition of a category in a neural region) (see also Dehaene-Lambertz et al., 2018, discussed below). However, both NR and GHS can be considered as instances of reading-based lateralized neuronal competition (or *reading-LNC*) theories (Rossion and Lochy, 2022), where local competition between word and face representations drive rightward face lateralization. Reading-LNC (see Figure 1 for a schematic) has been supported by multiple lines of evidence. First, compared to illiterate individuals, literate individuals show a reduction in face responses in the anatomical peak of the left VWFA (Dehaene et al., 2010). Second, the onset of reading instruction predicts both behavioral signatures of rightward face lateralization (Dundas et al., 2013) and lateralized neural responses to faces measured with EEG (Dundas et al., 2014). Similarly, assessing individual differences, Dundas et al. (2015) demonstrated that rightward laterality of the N170 to faces increased with a stronger LH N170 for words, and to a weaker extent, greater degree of right-handedness (among left- and right-handers). Lastly, computational modeling has supported the plausibility of the GHS claims (Plaut and Behrmann, 2011; Blauch et al., 2022a), demonstrating that the pressure for words to engage LH language representations can result in both leftward lateralization for words, and a compensatory weaker rightward lateralization for faces.

**Figure 1:**
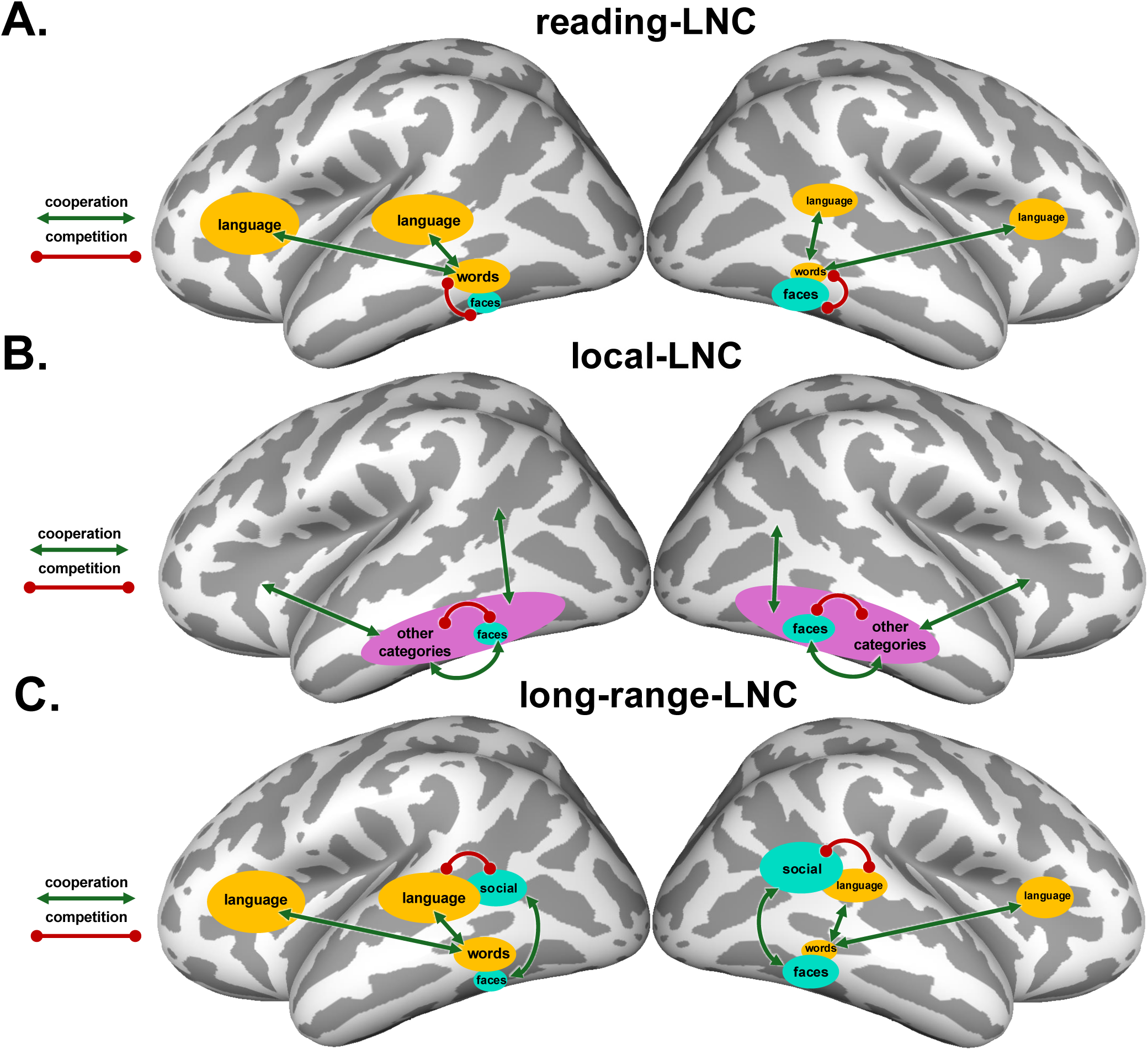
Visual schematic of the three lateralized neuronal competition (LNC) theories examined here. Competitive and cooperative relationships proposed in each theory are shown in individual panels. The size of blobs indicate the relative hemispheric specialization. We next describe these three theories, which are not mutually exclusive. **A**. *Reading-LNC* : learning to read gives rise to a left VTC laterality for words, to couple with language. Rightward VTC face laterality results from local competition with left-lateralized VTC word representations, which demand similar foveal inputs but require distinct representations from faces. **B**. *Local-LNC* : Representational overlap of different categories influences the sign and degree of correlated laterality. Rightward VTC face laterality emerges through local lateralized neuronal competition and cooperation more generally, not specific to words. Note: this theory does not specify the primary source(s) of lateralization in VTC, however, as shown with green cooperation arrows between "other categories" and other unspecified brain areas, it is presumed not to affect faces directly, but rather indirectly through local competition with representations that are affected by such external factors (as in reading-LNC, but more generically). **C**. *Long-range-LNC* : Language processing, which has an innate bias towards left lateralization, competes with social processing in the STSp, giving rise to rightward lateralization of social processing. VTC face laterality is induced by long-range coupling with social processing, similar to the long-range pressure of words to couple with language representations in both temporal and frontal areas.

## 1.4 Challenges to local competition between words and faces

Recently, however, there have been some important challenges to the claim that face lateralization is driven by local recycling or competition with emerging word representations. Two longitudinal studies have examined children’s reading acquisition to document the emergence of word selectivity, and to examine retroactively the site of eventual word-selective cortex before children were exposed to orthography. In the first study, Dehaene-Lambertz et al. (2018) found that word-selective cortex did not recycle previously face-selective cortex. Rather, eventual word-selective voxels appeared to show weak selectivity prior to the onset of reading, with some voxels showing mild preferences for tools, but not for faces. These findings led Dehaene-Lambertz and colleagues to revise the NR hypothesis, suggesting that word selectivity emerges from weakly selective voxels; in contrast, right-hemispheric face lateralization emerges due to a blocking mechanism, whereby left-lateralized word selectivity prevents the expansion of left-hemisphere face selectivity, yielding a relative rightward bias for faces. In a second study, Nordt et al. (2021) found that left-lateralized word selectivity emerged from voxels that were weakly selective for limbs but not body parts more generally (the Dehaene-Lambertz et al. (2018) study did not include a limb category). Limb selectivity was found to wane over development, with the more weakly limb-selective voxels converting to word-selectivity, along with other non-selective voxels. These findings led Kubota et al. (2023) to conclude that faces and words develop along independent trajectories (although this proposal leaves the rightward face lateralization unexplained). While the evidence does not support direct NR of face-selective cortex by words (Dehaene-Lambertz et al., 2018; Nordt et al., 2021), broader competitive relationships between reading and face processing would appear important in explaining the various empirical relationships found between reading skill and face lateralization (Ben-Shachar et al., 2011; Dehaene et al., 2010; Dundas et al., 2013, 2014; Kubota et al., 2019; Pinel et al., 2015).

Rossion and Lochy (2022) recently offered similar challenges to the proposed local competition between words and faces in VTC, noting inconsistency in the findings of studies that have attempted to assess this prediction. While Brederoo et al. (2020) found a relationship between word and face laterality measured behaviorally using reaction time, and Dundas et al. (2014, 2015) found a relationship between the laterality of word and face-related ERPs, fMRI studies examining individual differences have revealed a murkier picture with mixed findings but generally lacking support for a direct relationship between the laterality indices of word and face selectivity in VTC (Davies-Thompson et al., 2016; Pinel et al., 2015; Canário et al., 2020).

### 1.4.1 Long-range lateralized neuronal competition

Rossion and Lochy (2022) suggested an alternative mechanism of cortical competition, whereby competition between language processes (relevant for words) and spatial and social processes (more relevant for faces) outside of VTC results in a domain-relevant lateralization that then influences the lateralization of VTC, through a coupling mechanism similar to that proposed between visual word and language representations. While Rossion and Lochy (2022) referred to this theory as the *language-related lateralized neural competition* (LNC) theory, to contrast it with *reading-LNC* theories emphasizing the role of local, reading-induced competition within VTC, we will refer to it as the *long-range- LNC* (schematized in Figure 1). This emphasizes how VTC face lateralization depends on competition in areas that show long-range coupling with VTC, and contrasts it more precisely with the other theories examined in this work. Here, we focus on the possible role of social perception in influencing the laterality of face representations. Using multiple types of social interaction stimuli, Isik et al. (2017) demonstrated clustered selectivity for social interactions in the posterior superior temporal sulcus (STSp) of the right hemisphere. This selectivity was found to directly abut and partially overlap selectivity for dynamic face videos vs. dynamic videos of other object categories (Isik et al., 2017). Notably, even when analyzing the non-overlapping populations of voxels from each functionally-defined region, the region preferring social interactions retained a significant response to faces, and the region preferring faces retained a significant response to social interactions and multivariate information useful for decoding between positive and negative interactions. Social processing in the STSp has been studied extensively, revealing a complex map of selectivity for different subprocesses of social perception (McMahon et al., 2023; McMahon and Isik, 2023), with face perception considered one of several entry-points into higher-level social perception. However, rather than being entirely separable, the cortical selectivity profile suggest an intimate relationship between face perception and social perception in the STSp, one that might influence the laterality of the broader face processing network (Haxby et al., 2002).

The long-range LNC theory states that 1) language locally competes with some other function – likely social processing, but perhaps others – giving rise to a rightward laterality for that function, which 2) then has a downstream effect on the lateralization of face processing. The first claim has received recent empirical support, with demonstrations of a spatially precise competition between language and social processing in the STSp (in a social task reliant on spatial perception; Rajimehr et al., 2022), and a general complementary profile of RH lateralization for social processing across the LH dominant language network. Thus, this complementary rightward lateralization for social processing may encourage rightward lateralization of VTC face representations through long-range cooperative pressures between VTC and the STSp (see also Powell et al., 2018, for related ideas regarding coupling with medial prefrontal cortex playing a role in the location of VTC face selectivity). Intriguingly, Pinel et al. (2015)—who failed to find competition between word and face laterality in VTC—demonstrated a positive correlation between rightward VTC face laterality and reading skill (see also Dundas et al., 2013), as well as between both leftward VTC word and rightward face laterality with the leftward laterality of the superior temporal sulcus (STS) in a spoken language task. While this doesn’t address the coupling with social processing directly, the collection of findings is suggestive of a possible link.

### 1.4.2 General local lateralized neuronal competition

This final theory considers more general local competition within VTC between faces and other categories, without lending privileged status to words. Instead, this theory suggests that the entire graded topography of VTC is subject to lateralized competition and cooperation for representational and physical space (Figure 1). Accordingly, we term this theory *local-LNC*. This theory arises naturally from a locally interactive view of the topography of VTC, where topography emerges in a graded fashion due in part to the visual properties that must be learned in service of general high-level visual tasks (Blauch et al., 2022b; Doshi and Konkle, 2023; Prince et al., 2024). On this view, interactions between categories may be more generic, with hemispheric organization being one facet of topographic organization that is dependent on graded competition between domains. The graded competition may be initiated by coupling between a distal source and particular local representation, but the outcome for other categories depends on the similarity of local representations. Despite the appearance of local category processing modules, VTC is known to contain highly distributed information content (Haxby et al., 2001, 2011), where stimuli from non-preferred categories can be decoded from category selective areas. Such information appears to arise due to the systematicity of non-preferred responses, which is not idiosyncratic but extends across subjects (Downing et al., 2006; Prince et al., 2024) and is well captured by deep neural network models (Prince et al., 2024). This distributed information is thus not inconsistent with local functional specialization (Spiridon and Kanwisher, 2002), but it also points to the graded and interactive nature of representations in VTC. Critically, the overlap in neural responses for different categories predicts the behavioral ability to process multiple objects, with greater difficulty in processing multiple exemplars from categories with similar neural representations (faces and objects) than from categories with more distinct neural representations (faces and scenes) (Cohen et al., 2014). Similarly, neural representational similarity of pairs of categories in VTC strongly predicts visual search results (Cohen et al., 2017).

Thus, the trajectory of local lateralized competition may be determined largely by the similarity structure of VTC representations, not just the similarity of input and output demands. To the extent that some categories recruit lateralized processing outside of VTC, this could incite local lateralized competition with other categories depending on the overlap in representation. Notably, in addition to language lateralization impacting the lateralization of words, visual tool processing is known to elicit a left lateralized circuit in right-handed individuals (Johnson-Frey, 2004; Lewis, 2006)—including in VTC (Chao et al., 1999; Downing et al., 2006)—presumably due to visuomotor interactions supporting visually-guided tool use with the dominant (right) hand. Of interest, representations of tools are known to overlap specifically with representations of hands (Bracci et al., 2012; Knights et al., 2021), which are located next to general body representations (Bracci et al., 2010), themselves being next to face representations (Downing et al., 2001, 2006). Such systematicity may give rise to a relationship between face laterality and both tool and body laterality.

## 1.5 The present study

In the current work, we focus on elucidating the laterality of human ventral temporal cortex, with a focus on face processing. Since face representations begin to develop very early in life, longitudinal “recycling” studies (Nordt et al., 2021; Dehaene-Lambertz et al., 2018), which have been useful for studying word representations, may be less well suited to examine the emergence of face lateralization, since face representations do not "recycle" any pre-existing functions. Rather, assessing the variation across individuals may yield greater insight into general lateralized neuronal competition with face representations in VTC.

Here, with a sample of over 50 participants, we present the results of a neuroimaging study aimed at elucidating patterns of individual variability in the hemispheric organization of high-level visual representations. Additionally, we incorporate analyses of the large-scale Human Connectome Project (Barch et al., 2013; Glasser et al., 2013), power to test related ideas, benefitting both from the large number of subjects and the presence of additional tasks to probe brain activity. We begin by demonstrating strong individual variability in the topographic and hemispheric organization of VTC. We then use this variability as an inferential method to study the sources of lateralization in VTC, analyzing local competition and long-range coupling. Specifically, our data will be used to examine three related (non mutually-exclusive) theories regarding the role of lateralized neuronal competition (LNC) in the development of rightward face lateralization, extending the terminology of Rossion and Lochy (2022) (see Figure 1). Importantly, while the first theory proposes specifically *developmental* competition, the latter two theories could be implemented by competitive forces over development alone, or in combination with evolved changes in brain anatomy.

1. *Reading-LNC:* rightward VTC face laterality results from local competition with left-lateralized word representations;
2. *Local-LNC:* rightward VTC face laterality emerges through local lateralized neuronal competition and cooperation more generally, not specific to words, initiated by unspecified external cooperative pressures;
3. *Long-range-LNC:* VTC face and text laterality are induced by long-range coupling with external lateralized processes; specifically, VTC text laterality is induced by coupling with LH frontotemporal language areas, and VTC face laterality is induced by coupling with RH frontotemporal areas involved in processes that compete locally with language, such as social processing.

### 2 Methods

## 2.1 Main neuroimaging experiment

### 2.1.1 Participants

Individuals were recruited from the Carnegie Mellon University community, and were excluded from consideration they met any of the following criteria: left-handed, non-native English speaker, multilingualism in childhood, presence of metal in body, significant neurological or psychiatric history, claustrophobia, or any other contraindication for MRI. A total of 55 right-handed young adults (35 female, age mean=21.3, SD=3.89, Edinburgh handedness mean=86.9, SD=16.5) completed a 1-hour session of MRI scanning and two hours of behavioral experiments. Diffusion imaging and behavioral results were not analyzed for the results reported in this paper. An additional participant was recruited, but did not complete the imaging session and was thus excluded from analysis.

The procedures used in this study were reviewed and approved by the Institutional Review Board of Carnegie Mellon University. All participants gave informed consent prior to their participation, and were paid $50 for their participation.

### 2.1.2 Experimental protocol

Imaging data were acquired on a Siemens Prisma 3T scanner with a 64-channel head coil at the BRIDGE Center of Carnegie Mellon University and the University of Pittsburgh. The imaging sequence included a T1-weighted anatomical image, field-mapping image, diffusion-weighted images, and functional images. The anatomical image was collected first, followed by the field-mapping image, followed by two functional runs, followed by the diffusion-weighted images, followed by the final three functional runs. Thus, in total five runs of functional imaging data (TE=30 ms, TR=2000 ms) were collected, at isotropic resolution of (2mm)^3^. Each functional run lasted approximately 5 minutes, containing several mini-blocks of multiple images from a given category presented rapidly at fixation. We used a modified version of the functional localizer of (Stigliani et al., 2015), referred to as *fLoc*, but with 6 categories: words, faces, objects, inverted faces (or scrambled, see next section), inverted words, and non-word letter strings. The object, face, and inverted face categories contained two subcategories, cars and guitars for objects, and adult and children for faces, using the original stimuli from (Stigliani et al., 2015). Participants performed a 1-back stimulus identity task during the scan. They were told there would be some instances in which an exact image would repeat in two consecutive frames, and that they should press a button as quickly as possible to indicate detection of the stimulus repeat. A total of 24 repeats occurred per functional run, randomly interspersed across mini-blocks. Participants were given 1 second to press the button for their response to be counted as correct, and accuracy was reported at the end of each block to allow for monitoring of subject attention. Rest mini-blocks were intermixed with category mini-blocks, and were treated the same as stimulus mini-blocks in terms of frequency and duration. The order of mini-blocks was fully counterbalanced across categories, such that a mini-block of each category preceded a mini-block of each other category exactly once in a run. Each mini-block lasted 6s, containing 12 stimuli (or 0 in the rest condition), each presented for 400ms with a 100ms inter-stimulus interval (ISI). The first block began 10s after imaging acquisition, to allow for the scanner to reach a steady state. Additional imaging acquisition lasting 14s followed the final block of stimuli, allowing for the BOLD signal to return to baseline. Example stimuli and a schematic for the experimental design are shown in Figure 2.

**Figure 2:**
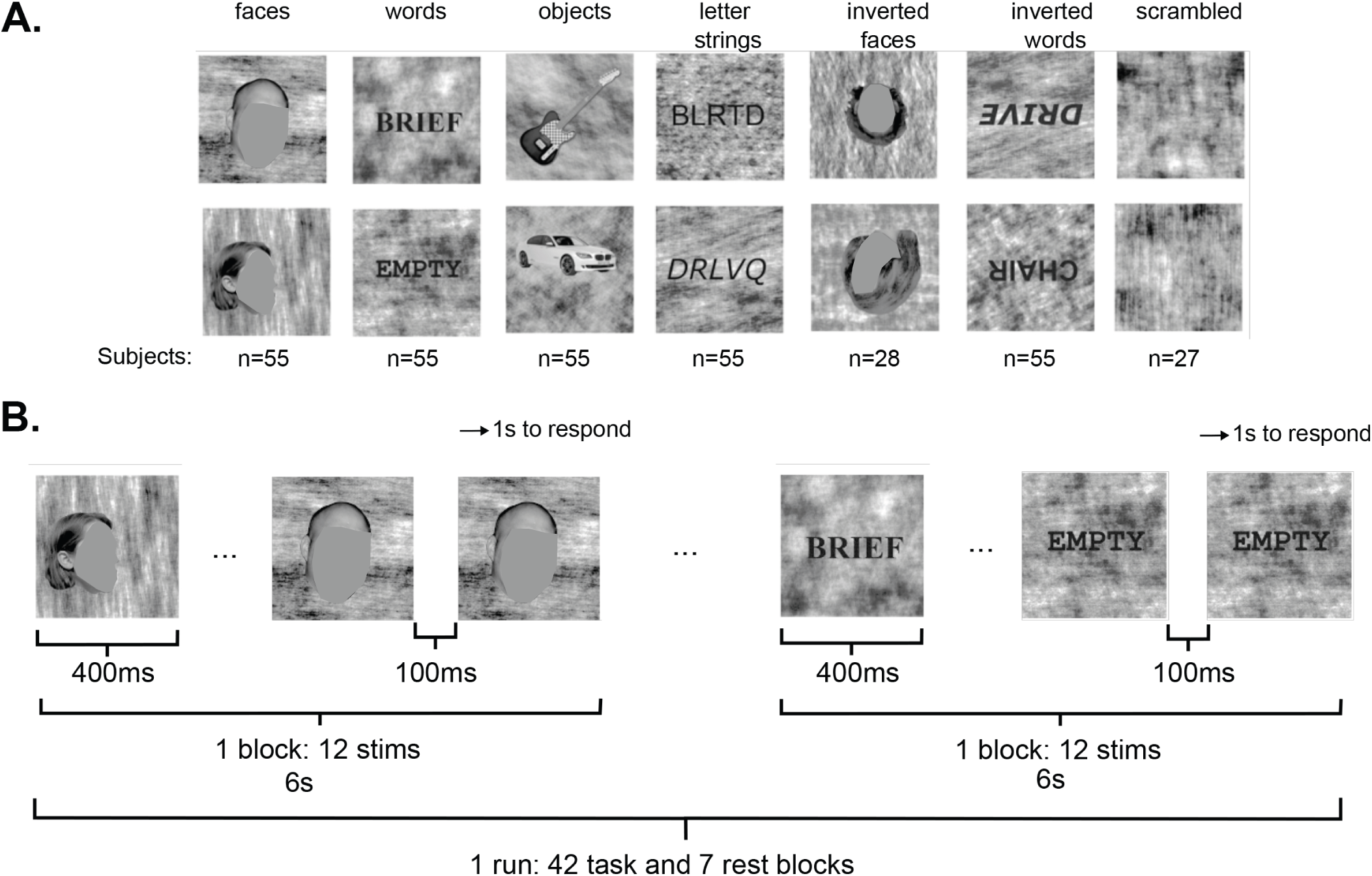
Experimental design. **A.** example stimuli from each of 6 categories, and the number of participants who viewed each category. **B.** a schematic of 1 run of the experiment. Note that the repeat condition could be anywhere within the block, not only at the end. Note: faces have been masked to conform to the policy of bioRxiv not to show human faces.

The data reported here were aggregated from two slightly different protocols. The first set of 28 participants performed the fLoc task with 6 categories: faces, inverted faces, words, inverted words, letter strings, objects (also used in Vin et al., 2024). The second set of 27 participants performed a similar fLoc task in the scanner, but the inverted faces condition was replaced with a scrambled image condition in order to assess selectivity relative to a baseline that allows for the removal of response estimation overlap in the raw beta coefficients, but does not enforce the selectivity map to contain only the most preferential responses for a given category. For the purposes of our individual difference analyses, and for any other analysis involved all subjects, the inverted faces and scrambled conditions are ignored so that identical contrasts can be constructed for each participant.

### 2.1.3 Functional imaging data pre-processing

The functional imaging data were preprocessed using *fMRIPrep* 1.4.1 (Esteban et al., 2018b,a), which is based on *Nipype* 1.2.0 (Gorgolewski et al., 2011, 2018). Many internal operations of *fMRIPrep* use *Nilearn* 0.5.2 (Abraham et al., 2014), mostly within the functional processing workflow. For more details of the pipeline, see the section corresponding to workflows in *fMRIPrep*’s documentation. Here, we quote the methods output directly from *fMRIPrep* as applied in our experiment, a procedure encouraged by the *fMRIPrep* authors to ensure accuracy of the methods description. Some modifications were made to remove unnecessary details regarding estimation of unused confounds, as well as to improve clarity:

For each of the 5 BOLD runs per participant, the following preprocessing was performed. First, a reference volume and its skull-stripped version were generated using a custom methodology of *fMRIPrep*. A deformation field to correct for susceptibility distortions was estimated based on two echo-planar imaging (EPI) references with opposing phase-encoding directions, using 3dQwarp (Cox and Hyde, 1997) (AFNI 20160207). Based on the estimated susceptibility distortion, an unwarped BOLD reference was calculated for a more accurate co-registration with the anatomical reference. The BOLD reference was then co-registered to the T1w reference using bbregister (FreeSurfer) which implements boundary-based registration (Greve and Fischl, 2009). Co-registration was configured with nine degrees of freedom to account for distortions remaining in the BOLD reference. Head-motion parameters with respect to the BOLD reference (transformation matrices, and six corresponding rotation and translation parameters) were estimated before any spatiotemporal filtering using mcflirt (FSL 5.0.9, Jenkinson et al., 2002). The BOLD time-series were resampled onto their original, native space by applying a single, composite transform to correct for head-motion and susceptibility distortions. Gridded (volumetric) resamplings were performed using antsApplyTransforms (ANTs), configured with Lanczos interpolation to minimize the smoothing effects of other kernels (Lanczos, 1964). These volumetric resampled BOLD time-series will be referred to as *preprocessed BOLD in original space*, or just *preprocessed BOLD*. The head-motion estimates calculated in the correction step were placed within confounds file for denoising in the general linear model (GLM), described later. Several additional confounding time-series were calculated based on the *preprocessed BOLD*: framewise displacement (FD), DVARS and three region-wise global signals. FD and DVARS were calculated for each functional run, both using their implementations in *Nipype* (following the definitions by Power et al., 2014). Additionally, a set of physiological regressors was extracted to allow for anatomical component-based noise correction (*aCompCor*, Behzadi et al., 2007). *aCompCor* was constrained within an anatomical mask designed

to be highly unlikely to contain relevant signal to the experiment. This mask was made up of the intersection of a subcortical mask, with the union of cerebro-spinal fluid (CSF) and white-matter (WM) masks. The subcortical mask was obtained by heavily eroding the brain mask, which ensures that it does not include cortical gray-matter (GM) regions. Masks were computed in T1w space, and projected into the native space of each functional run (using the inverse BOLD-to-T1w transformation) before performing the intersection and union across masks. Within the final mask, the *preprocessed BOLD* time-series was high-pass filtered using a discrete cosine filter with 128s cut-off before performing principal components analysis to yield the final aCompCor components, which were stored in the confounds file along with the other nuisance regressors.

### 2.1.4 fMRI General Linear Model (GLM)

fMRI data were analyzed using SPM12. For each participant, a general linear model was specified in order to model the effect of each stimulus category on the BOLD response of every voxel in the brain. Specifically, each mini-block was modeled as a single event specified by its onset and a duration of 6s. The design matrix thus contained 6 stimulus regressors, one per condition, each of which were convolved with a canonical hemodynamic response function. A reduced set of fMRIPrep-generated confounds was retained for nuisance regression; specifically, we retained 6 motion parameters (X, Y, Z motion and rotation), the top 6 principal components of the aCompCor decomposition, and the framewise-displacement, yielding a total of 13 nuisance regressors, along with a runwise mean regressor, which were appended to the design matrix. Finally, an autoregressive-1 model was used within SPM12 to reduce the effects of serial correlations. We fit voxels only within the brain mask defined by *fMRIPrep*. Estimation of the GLM resulted in a beta-weight for each stimulus condition and run, which were used for subsequent univariate and multivariate analyses.

### 2.1.5 Regions of interest

The primary region of interest (ROI) for this work is a large ventral temporal cortical (VTC) area, whose anatomical mask consisted of the union of the inferior-temporal and fusiform ROIs defined in the Freesurfer *aparc* atlas. We also defined multiple regions of interest outside of VTC, using the Glasser atlas (Glasser et al., 2016), following the approach of (Vin et al., 2024). Inferior frontal gyrus (IFG) was defined as the union of areas ‘44’,‘IFJa’,‘6r’, and ‘IFSp’; superior temporal sulcus and gyrus (STSG) was defined as the union of areas ‘STSdp’,‘TPOJ1’,‘A5’, and ‘STV’; precentral gyrus was defined as the union of areas ‘55b’ and ‘PEF’; IFG pars orbitalis (IFGorb) was defined as the union of areas ‘IFSa’, and ‘45’; Intraparietal sulcus (IPS) was defined as the union of areas ‘IP0’, ‘IP1’, ‘IP2’, and ‘IPS1’; early visual cortex (EVC) was defined as the union of areas ‘V1’, ‘V2’, and ‘V3’.

### 2.1.6 Calculation of fMRI hemispheric selectivity and laterality index

Hemispheric selectivity for various contrasts was computed separately and used to compute the laterality index (LI). The intersection of two masks—anatomical and stimulus-responsive—was used to constrain voxel selection for the computation of selectivity in each hemisphere. Each ROI was associated with one or more anatomical parcels used to construct the anatomical mask. The stimulus-responsive mask was defined as voxels whose response was significantly greater during the stimulus periods than rest periods, with *p <* 10*^−^*^4^. Selectivity was computed from whole-brain statistical maps computed in SPM12, which used contrasts comparing voxelwise activation of some set of conditions to another. As words, letter strings, and inverted words evinced similar responses in ventral regions (see Vin et al., 2024, for explicit consideration of differences in activation across text conditions within this dataset), we grouped words, inverted words, and letter strings into a “text” domain. Responses to inverted faces were not analyzed in this work. Contrasts were balanced in total weight across numerator and denominator, as well as across any sampled domains within the numerator or denominator (i.e. faces, objects, text), to avoid biasing our selectivity metrics to the more heavily sampled domains (i.e. text). We computed *text selectivity* with the contrast (Words + Inverted Words + Letter Strings)/3 *−* (Faces + Objects)/2. Likewise, to compute *face selectivity*, we used the contrast (Faces) *−*((Words + Inverted Words + Letter Strings)/3 + Objects)/2. Last, we computed object selectivity as Objects *−* (Faces + (Words + Inverted Words + Letter Strings)/3)/2. We refer to these contrasts as the **full baseline** contrasts, indicating that the baseline contains all of the domains, as typically done in fMRI studies. In the next section and in Table 1, we explain our use of alternative contrasts with different baselines. We examined three dependent measures of hemispheric selectivity: summed positive t-statistics, peak t-statistic, and number of selective voxels. Where not stated, we used summed selectivity as the metric of selectivity, as it best captures the total profile of selectivity considering both the size of the region as well as the magnitude of the selectivity within the selective region; while it is useful to correct for total cortical volume of the bilateral anatomical ROI when assessing hemispheric selectivity, the laterality index is unaffected by this correction. An illustration of our approach can be seen in Figure 3. For each metric, selectivity was computed for a given category *C* in selected voxels of a given hemisphere *H*, denoted *S_C,H_* . Finally, the laterality index for a given category *C*, *LI_C_*, was computed as:

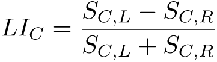

**Figure 3:**
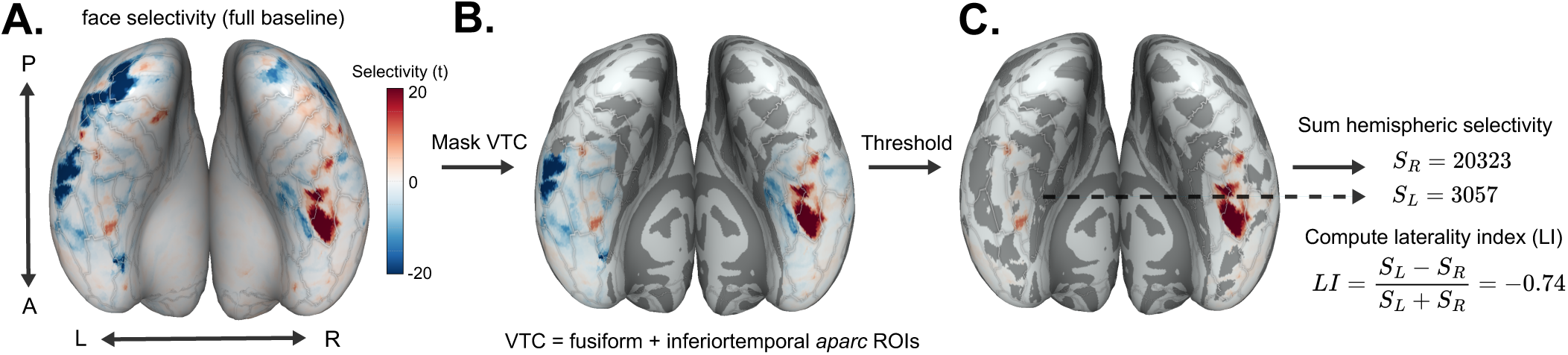
Computing summed hemispheric selectivity and laterality index (LI) in VTC, using face selectivity in an example participant. The ventral inflated cortical surface is shown. Posterior-anterior axes are labeled with *P* and *A*, and left and right hemispheres with *L* and *R*. The t-statistic map (left) is first masked anatomically by the VTC ROI, and then thresholded at 0. Positive t-statistics are then summed to compute selectivity in the left and right hemisphere, *S_L_*and *S_R_*, respectively. Finally, the laterality index is computed.

**Figure 4:**
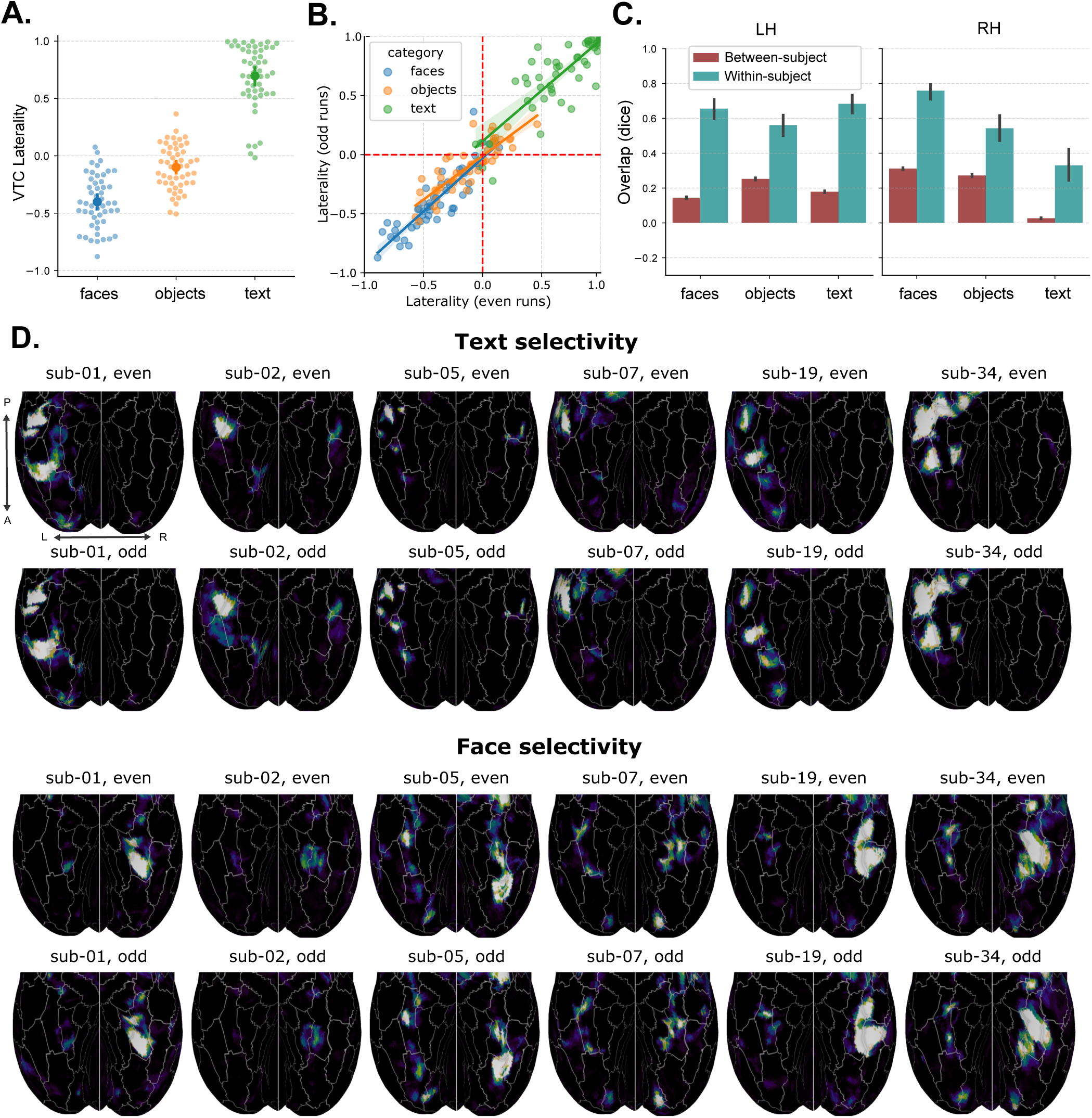
Individual differences in VTC laterality. **A.** Distributions of LIs for each category contrast. **B.** Consistency of individual LIs across even and odd subsets of runs. **C.** Overlap of category-selective areas across even and odd runs, across-participant (blue), and within-participant (orange). Overlap is computed as the dice coefficient applied to category-selective masks computed from even or odd runs, thresholded at *p <* 0.001. **D.** Example individual participant maps for face and text selectivity, demonstrating large inter-individual variability and within-individual reliability.

**Table 1:** Functional contrasts used in each pairwise LI comparison of our in-house experiment (Figure 5). Where *β_i,F_* , *β_i,O_*, *β_i,W_* , *β_i,L_*, *β_i,I_* are the beta coefficients for faces, objects, words, letter strings, and inverted words, respectively, for a given run number *i*, we let 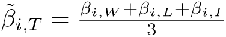 . Note that the "fixation baseline" uses an implicit baseline, since the GLM-derived beta coefficients are inherently in reference to the unmodeled fixation baseline in the experimental design. Where not otherwise stated in the main text, the "full" baseline serves as the default.

| Category 1 | Category 2 | Baseline | Contrast 1 | Contrast 2 |
| --- | --- | --- | --- | --- |
| Faces | Text | Full | $\sum_i^{1,3,5} \beta_{i,F} - \frac{\tilde{\beta}_{i,T} + \beta_{i,O}}{2}$ | $\sum_i^{2,4} \tilde{\beta}_{i,T} - \frac{\beta_{i,F} + \beta_{i,O}}{2}$ |
| Faces | Text | Held-out (Objects) | $\sum_i^{1,3,5} \beta_{i,F} - \beta_{i,O}$ | $\sum_i^{2,4} \tilde{\beta}_{i,T} - \beta_{i,O}$ |
| Faces | Text | Fixation | $\sum_i^{1,3,5} \beta_{i,F}$ | $\sum_i^{2,4} \tilde{\beta}_{i,T}$ |
| Objects | Text | Full | $\sum_i^{1,3,5} \beta_{i,O} - \frac{\tilde{\beta}_{i,T} + \beta_{i,F}}{2}$ | $\sum_i^{2,4} \tilde{\beta}_{i,T} - \frac{\beta_{i,O} + \beta_{i,F}}{2}$ |
| Objects | Text | Held-out (Faces) | $\sum_i^{1,3,5} \beta_{i,O} - \beta_{i,F}$ | $\sum_i^{2,4} \tilde{\beta}_{i,T} - \beta_{i,F}$ |
| Objects | Text | Fixation | $\sum_i^{1,3,5} \beta_{i,O}$ | $\sum_i^{2,4} \tilde{\beta}_{i,T}$ |
| Faces | Objects | Full | $\sum_i^{1,3,5} \beta_{i,F} - \frac{\beta_{i,O} + \tilde{\beta}_{i,T}}{2}$ | $\sum_i^{2,4} \beta_{i,O} - \frac{\beta_{i,F} + \tilde{\beta}_{i,T}}{2}$ |
| Faces | Objects | Held-out (Text) | $\sum_i^{1,3,5} \beta_{i,F} - \tilde{\beta}_{i,T}$ | $\sum_i^{2,4} \beta_{i,O} - \tilde{\beta}_{i,T}$ |
| Faces | Objects | Fixation | $\sum_i^{1,3,5} \beta_{i,F}$ | $\sum_i^{2,4} \beta_{i,O}$ |

### 2.1.7 Functional contrasts

To assess competition across domains, we used multiple different functional contrasts, built from beta coefficients derived from a single underlying GLM. The purpose of this approach was to minimize confounding factors in the establishment of correlations across domains. In all cases, statistical confounding was prevented by using independent runs of data to compare LI values across domains; however, category-based confounding may still persist by way of different selectivity contrasts containing overlapping sets of categories. Considering the standard full baseline for faces and text (defined in the previous section), it is clear that both positive and negative biases exist in the correlation of the two contrasts. First, to the extent that object responses and LI are reliable across subsets of runs, the presence of objects in both denominators induces a positive correlation between the two contrasts. Perhaps more problematically, under the same assumption of reliability for faces and text, the swapped presence of faces and text in numerator and denominator across the two contrasts induces a bias towards negative correlation. Thus, comparisons using full contrasts may be inappropriate for discovering true patterns of correlation across participants, being biased by category-based confounds due to overlap in the contrasts. The simplest solution is to assess the LI of each domain against the fixation baseline, using a one-sample t-test of beta coefficients against 0. However, this may fail to eliminate a large degree of shared variability that may be overestimated due to the fast event-related design used here. Thus, in addition to examining the 1) full baseline and 2) fixation baseline, we additionally examine 3) the **held-out domain baseline**, where in the comparison of LI for two domains (e.g. face and text), the third domain is used as a common baseline (i.e. faces vs. objects, text vs. objects), and 4) the fixation baseline with the LI of the held-out category regressed out of the LI of each domain in the comparison. These four approaches make up the four columns of Figure 5, and are described explicitly in Table 1 below

**Figure 5:**
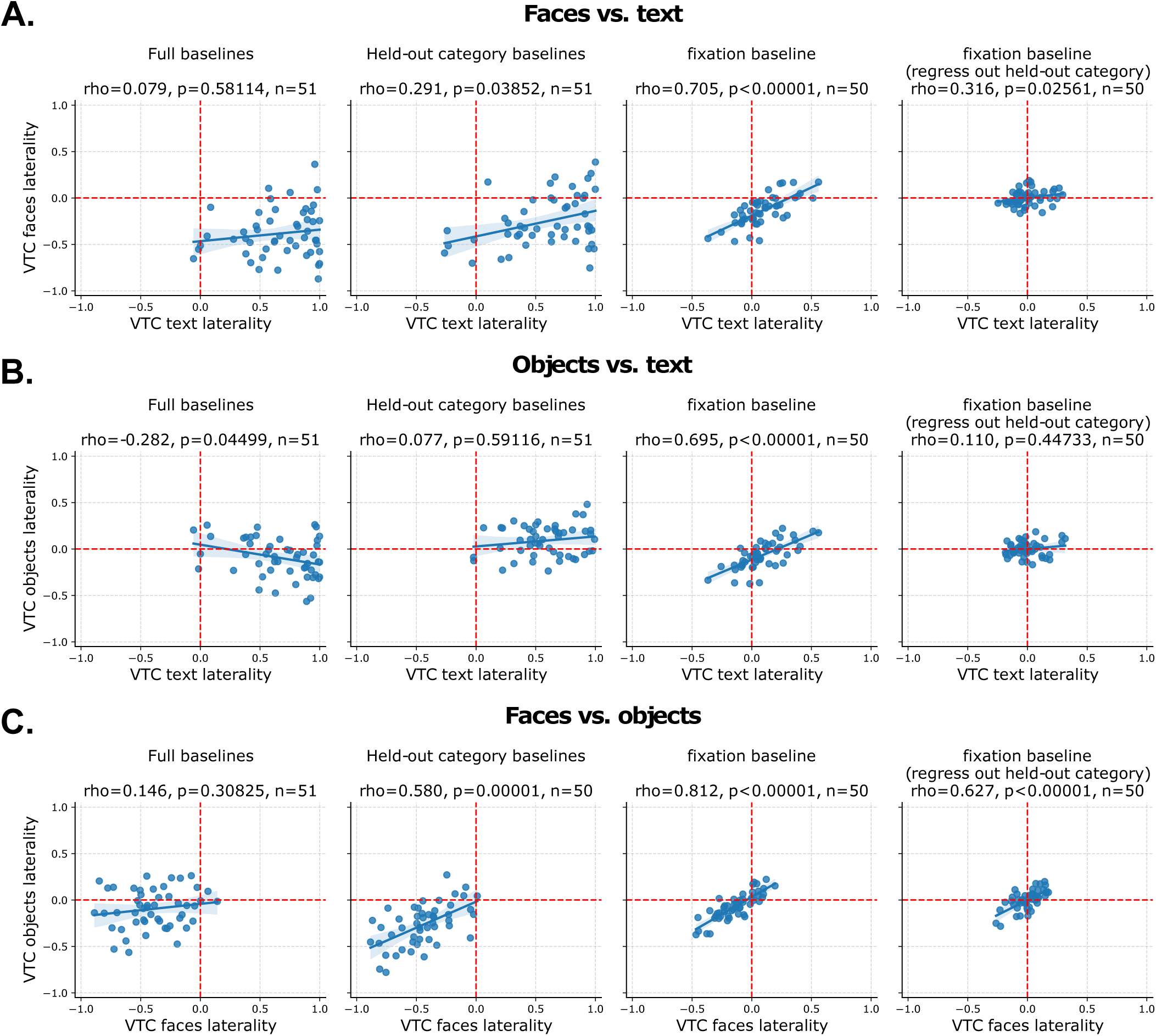
Pairwise laterality index (LI) comparisons for faces, objects, and text. **A.** face vs. text selectivity. **B.** object vs. text selectivity. **C.** face vs. object selectivity. For each subfigure, the first (leftmost) plot uses the full baseline (all categories), the second plot uses the held-out domain as the baseline, the third plot uses a fixation baseline, and the fourth plot uses a fixation baseline after regressing out the held-out category’s LI (using selectivity computed against a fixation baseline) from both LI values. In each case, even runs are used to compute the LI for the X-axis, and odd runs are used to compute the LI for the Y-axis. For full details on contrasts, see Table 1.

### 2.1.8 Whole-brain atlas-based laterality

In some analyses, we computed LI across the whole-brain using a fine-grained cortical parcellation. For this purpose, we used the Yan homotopic atlas with 500 parcels per hemisphere (Yan et al., 2023).

### 2.1.9 Cortical surface visualization

Pycortex (Gao et al., 2015) was used for visualization. All visualizations were performed in *fsaverage* surface-space, following a sulcal-based alignment (Fischl et al., 1999) implemented in Freesurfer. The boundaries of parcels from the Human Connectome Multi-Modal Parcellation (HCPMMP) atlas (Glasser et al., 2016) were manually drawn in Inkscape and added to the Pycortex ‘overlays.svg‘ file as a visual guide to viewing the surface images.

## 2.2 Analysis of the Human Connectome Project

In addition to the analyses of our in-house dataset, we took advantage of the large-scale Human Connectome Project (HCP) dataset to perform additional, complementary analyses. For computational convenience, we downloaded pre-computed category selective maps available on Amazon Web Services S3 storage, under the HCP_1200 folder. We focused on the Working Memory, Language, Social, and Emotion tasks. These tasks have been discussed extensively in prior literature, so we will discuss only the basics here and refer the reader to Glasser et al. (2016) for more specific details.

We extracted surface-based contrast maps in the *32k_fs_LR* group CIFTI space of the HCP preprocessing pipeline (Glasser et al., 2016), using standard sulcal-based alignment rather than the multimodal-surface-matching procedure (Robinson et al., 2014), due to the latter’s use of functional data to compute the alignment, which could

introduce dependencies between estimates of different functional topographies. We acquired the maps smoothed with a Gaussian kernel of 2mm FWHM, which was the smallest amount of smoothing available. The hcp_utils(https://rmldj.github.io/hcp-utils/) Python package was used as a source of a CIFTI-based HCPMMP atlas, similar to the one used in our main analyses. The HCPMMP atlas was used for all selectivity analyses. We constructed a VTC ROI as the union of several HCPMMP parcels: V8, pIT, FFC, VVC, PHA1, PHA2, PHA3, TE2p, TF, PH, VMV1, VMV2, VMV3. This VTC ROI—shown in the inflated surface panels of Figure 6—is similar to the VTC ROI used in the main experiment (see Figure 3). Selectivity analyses were performed in the *32k_fs_LR* space; however, for the purposes of visualization in PyCortex (Gao et al., 2015), we transformed relevant maps to the standard higher resolution*fsaverage* cortical surface space using Connectome Workbench tools wrapped within the hcp_utils package.

**Figure 6:**
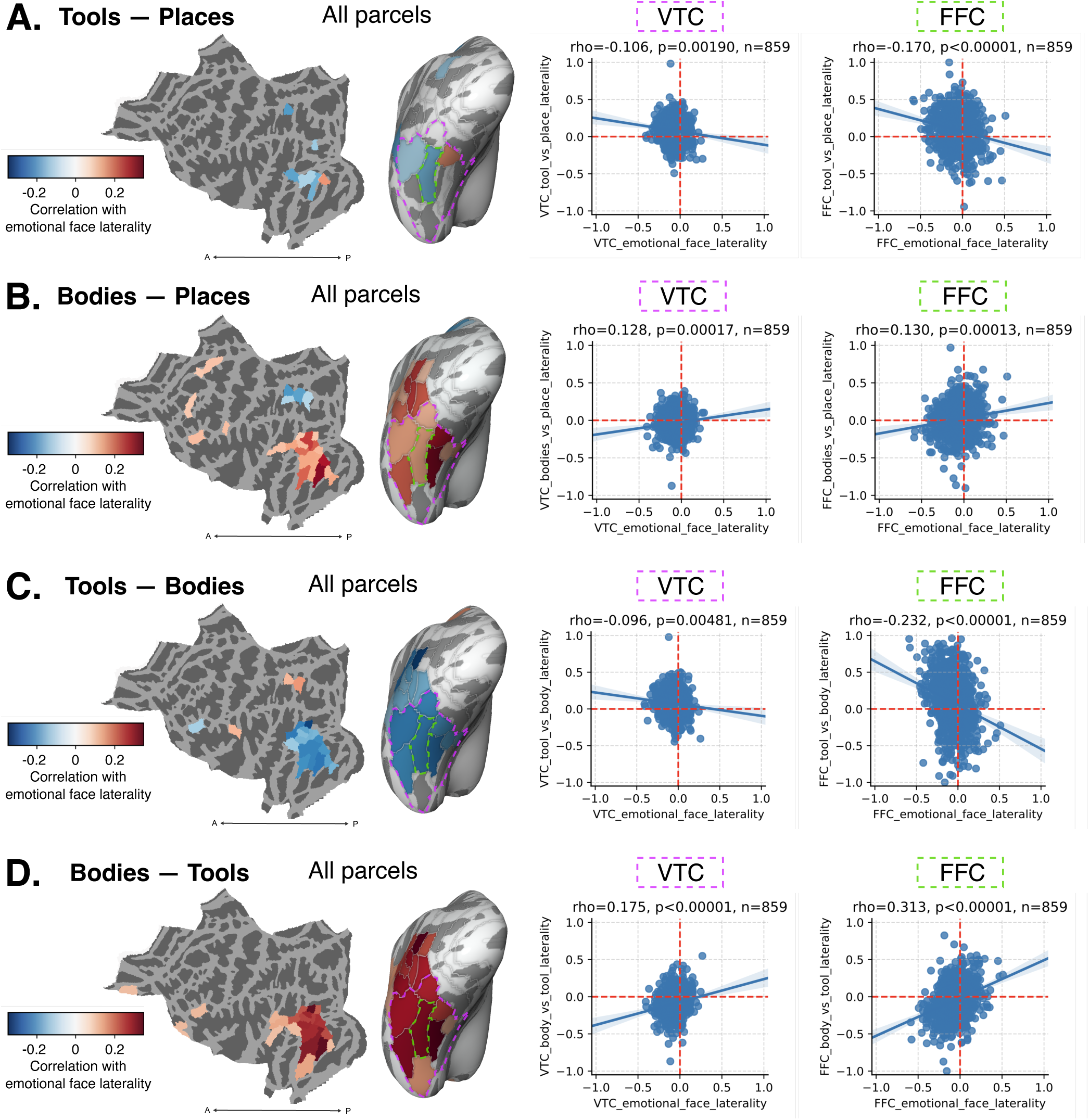
Comparing the laterality index (LI) for the faces*−*shapes contrast in the Emotion task, with LI for other contrasts encompassing tools and bodies. **A-D.** Left: parcel-level correlation, computed within each parcel of the HCPMMP atlas. Significant correlations following FDR correction over 180 regions are plotted in a flat map and inflated ventral view. Right: correlations for individual regions, including ventral temporal cortex (VTC; pink outline), a large region made up of several HCPMMP parcels and the fusiform face complex (FFC; green outline). **A.** Tools*−*places. **B.** Bodies*−*places. **C.** Tools*−*bodies. **D.** Bodies*−*tools.

### 2.2.1 Visual working memory task

The visual working memory (WM) task consisted of a block design of visual categories—faces, bodies, tools, and places—with a 0-back or 2-back working-memory task, which was used to index category selectivity in VTC and neural mechanisms for visual working memory. Our focus is on the former usage. For the WM experiment, HCP provides contrasts of each category against baseline (fixation), as well as against the average of all other categories. The former contrast essentially accounts for the signal in beta weights for a given category across runs, given the noise in the estimates, whereas the latter contrast is a more typical index of “selectivity” for one category versus the others. Neither of these choices is ideal for assessing individual differences in selectivity across participants: contrasting each category versus baseline is too lenient and results in widespread positive correlations between all conditions, whereas contrasting each category versus the average of all others introduces negative correlations due to statistical dependence between the contrasts for each category. We constructed new contrasts by making use of the category versus baseline contrasts, which were constructed across two experimental runs. We chose “places” as a baseline category to compare bodies, tools, and faces. These contrasts were constructed by subtracting the places versus fixation contrasts from each category versus fixation contrast.

### 2.2.2 Language task

The language (LANG) task consisted of a block design of story listening or math problem-solving blocks (Binder et al., 2011; Glasser et al., 2016). The contrast of primary interest was stories versus baseline, following Glasser et al. (2016) and Rajimehr et al. (2022).

### 2.2.3 Social task

The social (SOCIAL) experiment consists of a block design alternating blocks of interactions between simple shapes that are either social or random in nature, with the task being to determine whether the interaction is social or not (Glasser et al., 2016). As in Rajimehr et al. (2022), we used the “theory of mind (TOM) *−* random” contrast as an index of social processing.

### 2.2.4 Emotion task

The emotion (EMOTION) task presents blocks of emotional face stimuli alternating with blocks of non-emotional simple shape stimuli (e.g., rectangles, ovals). Participants are tasked with matching a left or right stimulus with a target stimulus above the two sample stimuli. We computed emotional face selectivity and LI using the contrast (faces *−* shapes).

### 2.2.5 Whole-brain atlas-based laterality

As in the main experiment, we performed an analysis based on the LI computed in a fine-grained cortical parcellation. For greater facilitation with the CIFTI-based HCP data, we used the HCP parcellation, containing 180 parcels per hemisphere (Glasser et al., 2016).

### 2.2.6 Social and language processing in the homotopic language network

To assess the long-range LNC theory, we used a recently developed parcellation of language network regions used to demonstrate competition between social and language processing (Rajimehr et al., 2022), based on group-level language activation in the HCP task. 8 regions were parcellated: "Broca" in the inferior frontal gyrus, along with "55b", "perisylvian language area (PSL)", "parietal area G inferior (PGi)", "posterior superior temporal sulcus (STSp)", "anterior superior temporal sulcus (aSTS)", "anterior superior temporal gyrus (STGa)", and "superior frontal lobule (SFL)", all of which were named based on the most closely related HCP anatomical parcels, which were otherwise not used in areal definition. These regions are shown in Figure 8A and are described in more detail in (Rajimehr et al., 2022).

**Figure 7:**
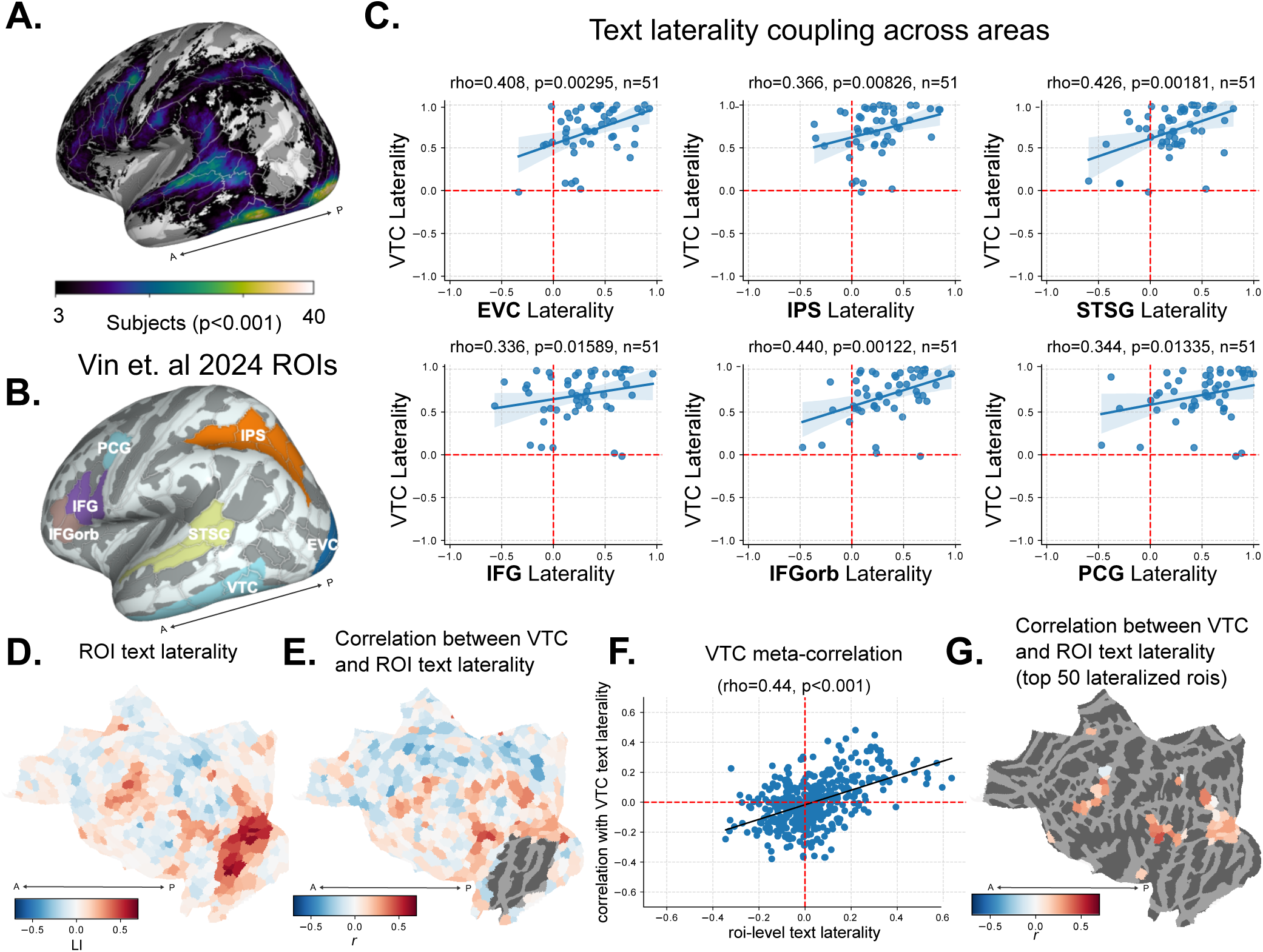
Laterality of extra-visual areas correlates with the hemispheric profile of VTC text selectivity. **A.** Text selectivity across the cortical surface (# of participants with significant activation, p<0.001). **B.** Anatomical parcels chosen for laterality comparisons, consisting of language network, dorsal visual, and early visual areas, along with VTC. **C.** Laterality of VTC text selectivity is correlated with laterality of text selectivity across this distributed network. Note that for **C.** and **E-F** below, separate sets of runs were used to compute laterality in VTC and other regions, eliminating trial-level correlations. **D.** Mean of individual atlas parcels across participants. Red corresponds to leftward lateralization, whereas blue corresponds to rightward lateralization. **E.** Correlation of individual patterns of laterality in each parcel with that of VTC, across participants, excluding parcels that overlap with the large VTC parcel. **F.** Scatter plot comparing the parcel-level mean laterality with the across-participant correlation of laterality between the parcel and VTC, along with a line of best fit and result of a spearman correlation across parcels. **G.** Results of **B.** masked to show only the 50 parcels with the largest mean laterality. The vast majority of these lateralized parcels show an across-participant correlation in laterality with VTC.

**Figure 8:**
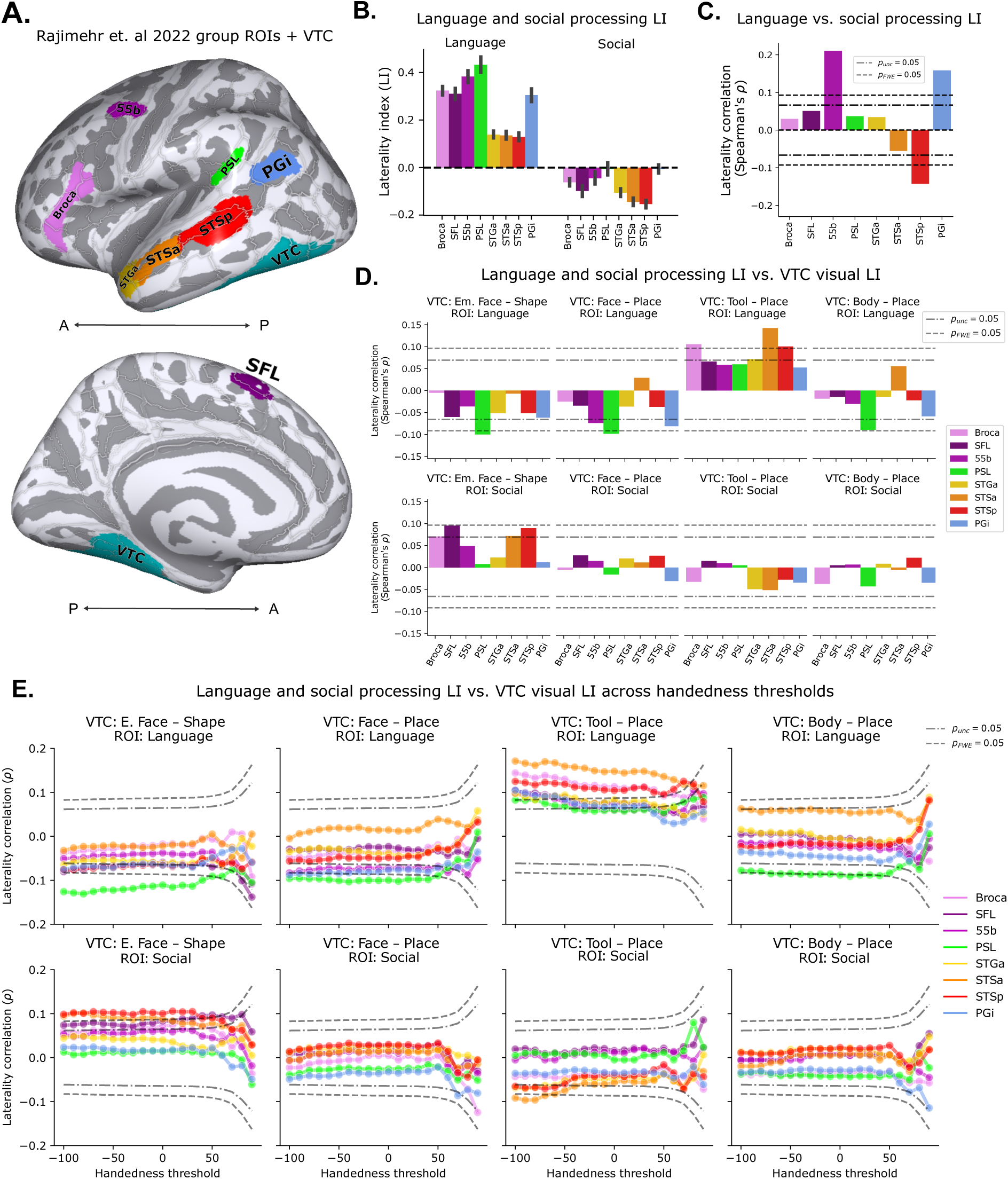
Long-range coupling in laterality for high-level vision, and language and social processing within functionally defined language ROIs. **A.** Group-level language parcels defined by (Rajimehr et al., 2022), plotted on the lateral and medial *fsaverage* surfaces along with the VTC parcel. **B.** Laterality index (LI) of language and social processing computed within each parcel. Here and in the following panels, we used the "story" contrast for language processing, and the "theory of mind (TOM) - random" contrast for social processing, as in (Rajimehr et al., 2022). **C.** Inter-subject correlation of language and social LI within each parcel. **D.** Inter-subject correlation of language and social LI in language parcels with visual processing LI in VTC, restricting to right-handed individuals (Edinburgh handedness > 40). **E.** Same as **D.**, but computed across all handedness thresholds; a threshold of 0 corresponds to including all subjects with a numerical right handedness, while a threshold of -100 corresponds to including all subjects including all strongly left-handed subjects.

#### 3 Results

## 3.1 Individual differences in VTC organization

Our first goal was to determine the degree of individual variation present in VTC organization by gathering and analyzing our own empirical data. Figure 4A shows the resulting distribution of laterality indices (LIs) for each category contrast, demonstrating that there is substantial variation in the magnitude of laterality for each domain (even among the right-handed college-aged population examined in this study). At the group-level, as expected, face selectivity is significantly right-lateralized and text selectivity is significantly left-lateralized, while object selectivity is weakly right-lateralized; group-level maps are additionally plotted in Figure S1.

We next quantified the reliability in subject-level LIs, by correlating LI values across two independent splits of the data (using either the even or odd runs) from the same participant, with the relationships shown in Figure 4B. We found that the reliability in LI is extremely high for each category (all Spearman’s *ρ >* 0.8, all *p <* 10*^−^*^10^).

These reliable within-individual differences can be further visualized in Figure 4D, which shows example individual participant maps for face and text selectivity (all subjects are shown in Figures S6 and S7. In addition to stable individual differences in LI, differences in within-hemisphere topography—including the location, size, and number of selective regions—are clearly present.

To quantify the general variability in topography across participants, and its reliability within participants, we computed the overlap of category-selective regions across participants and compared it with the overlap across independent splits of the data within participants (Figure 4C). Specifically, we binarized the *fsaverage* selectivity maps to be 1 if positive selectivity exceeded p<0.001, and 0 otherwise; we then computed the dice coefficient across maps masked within the VTC region. Whereas the overlap within participants is relatively high for most regions, indicating reliability, overlap is much weaker across participants, indicating the presence of substantial individual differences in functional topography.

Having established the presence of individual differences in VTC organization, we next sought to examine how different aspects of VTC organization are related to each other (across participants), in order to draw implications for the various theories of functional competition.

## 3.2 Local competition between faces and words

Rossion and Lochy (2022) argued that a key prediction of reading-LNC theories like NR and GHS is that the lateralization of faces and words should be linked or interdependent within VTC. We tested this prediction in young adults with competent reading skills. We compared laterality of faces, text, and objects, pairwise across participants, using multiple fMRI contrasts (see section 2.1.7). In all cases, we used separate subsets of runs to compare laterality to ensure statistical independence of the contrasts. We first assessed the relationship between face and text LI, shown in Figure 5A, which these two accounts predict should be negatively correlated. In contrast to this prediction, we found no clear correlation between LIs for faces and text when using the full baseline (*r* = 0.079, *p >* 0.05). When using responses to the held-out category (objects) as the baseline, we found a small positive correlation in face and text LI (*r* = 0.079, *p >* 0.05). Lastly, when using a fixation baseline^1^ (i.e. computing selectivity as a one-sample t-test of beta coefficients against 0), we found a strong positive correlation (*r* = 0.705, *p <* 0.00001). This correlation is largely reduced, but not abolished, when regressing out object versus fixation LI from both face and text LIs (*r* = 0.316, *p <* 0.05). In contrast to predictions from reading-LNC theories, rather than being negatively correlated, the LIs of faces and text appear to be positively correlated, except when using the full baseline for which there is a category-based bias towards negative correlation.

Importantly, the lack of evidence for lateralized competition between words and faces was not changed by using two more typical measures of selectivity indexing either the strength (peak) or size (number of selective voxels) of the selective region (Figures S4, S5). Taken together, these results do not provide evidence supporting the reading-LNC (NR and GHS) hypothesis that VTC face laterality emerges from local competition with word representations (Rossion and Lochy, 2022; Behrmann and Plaut, 2020; Dehaene et al., 2010).

As noted above, prior work has suggested that words may compete more broadly with object-responsive cortex, rather than with face-responsive cortex specifically (Kubota et al., 2019). Accordingly, we also compared the LI of objects and text across participants (Figure 5B). Using the full baseline, we found a small negative correlation (*r* = *−*0.282, *p <* 0.05), indicative of competition. However, when the held-out category of faces alone is used as a baseline in selectivity contrasts, we found a lack of competition (*r* = 0.077, *p >* 0.05), indicating that the prior result was likely due to the presence of the category-based confound. When using the fixation baseline, there is a very strong positive correlation between object and text LI (*r* = 0.695, *p <* 00001); however, this correlation is abolished when regressing out the face versus fixation LI (*r* = 0.11, *p >* 0.05).

## 3.3 Local correlation between faces and objects

Prior work demonstrated that the overlap in neural representations for faces and objects predicted the behavioral cost in representing exemplars from each of these categories, which was stronger than that for more distinct categories (e.g. faces and scenes) (Cohen et al., 2014). To determine whether there is also an overlapping *laterality* for faces and objects, we next compared their laterality patterns across participants (Figure 5C). Using the full baseline for each, we found a weak and non-significant trend towards positive correlation in LI (*r* = 0.146, *p >* 0.05). However, using the held-out text baseline, we found a strong correlation in face and object LI (*r* = 0.58, *p <* 0.0001). As both the lack of positive correlation in the full baseline, and the strong positive correlation in the held-out baseline, could be driven by category-based confounding in the baselines, we next assessed the results using the fixation baseline. As in the previous cases, there is a strong positive correlation in LI (*r* = 0.812, *p <* 0.00001). As a final test, we examined this correlation after regressing out text vs. fixation LI; the correlation remained strong (*r* = 0.627, *p <* 0.00001). This suggests that faces and objects exhibit a true correlation in their laterality across participants, despite faces being typically more lateralized, in line with the local-LNC theory.

## 3.4 Local interactions between faces and other categories

We next evaluated the claim of the local-LNC theory that generic local competition within VTC, not specific to words or reading, shapes the lateralization of face processing. For this purpose, we leveraged the Working Memory (WM) experiment of the Human Connectome Project (HCP), which has the advantage of having a much larger cohort of participants and thus greater statistical power to detect any effects. As discussed in Methods 2.2, this experiment consists of blocks of faces, places, bodies, and tools. Noting that the selectivity for places was substantially more medial than that of faces, bodies and tools, we opted to use places as a common baseline. Group-level category-selective maps of standard and modified contrasts can be seen in Figure S2. Faces, while present in the WM task, were also shown to participants in a separate Emotion experiment, consisting of a block design of emotional faces, and non-emotional simple shape stimuli. This yields the opportunity for an independent comparison of face laterality index (LI) and LI for other categories, not confounded by shared or opposing selectivity baselines. As a sanity check, we confirmed that face LI is similar across WM and Emotion tasks, throughout VTC (*r* = 0.483, *p <* 0.00001), and particularly in the fusiform face complex (FFC) parcel (*r* = 0.683, *p <* 0.00001). Thus, emotional face LI is a suitable independent measure of face LI to compare with body and tool LI from the WM experiment. We used contrasts of bodies*−*places, tools*−*places, and emotional faces*−*shapes to compute summed selectivity in each parcel of the symmetric HCPMMP atlas. Lastly, we computed LI values for each pair of homologous parcels.

To assess lateralized competition between face representations and those of bodies and tools, we correlated face LI with body and tool LI for each parcel, as well as at the level of the full VTC. For tools*−*places LI (Figure 6A), we found significant negative correlations with face LI within VTC, notably in the FFC parcel (*ρ* = 0.17, *p <* 0.00001). For bodies*−*places LI (Figure 6B), we found significant *positive* correlations with face LI within VTC, including in FFC (*ρ* = 0.13*, p <* 0.001), along with a broader swath of individual regions across ventral and lateral occipitotemporal cortex, peaking in the parcel immediately medial to FFC, known as the ventral visual complex (VVC; *ρ* = 0.316, *p <* 0.00001). These results are in line with the idea of local representational cooperation between faces and bodies, and local representational competition between faces and tools.

The correlations between the laterality indices of tools/bodies and faces are largely opposite in sign and might be complementary. To test this, we computed the contrasts tools*−*bodies and bodies*−*tools, and performed the same analyses. The tools*−*bodies contrast (Figure 6C) reveals a larger number of negative correlations in VTC (despite a weaker correlation at the level of the full VTC swath), and an increase in the magnitude of the negative correlation with emotional face LI in FFC. Similarly, bodies*−*tools (Figure 6D), reveals a strengthening of positive correlations with emotional face LI, including increases in VTC and FFC. Notably, the magnitude of the correlation is strongest in VTC and FFC when using the contrast bodies*−*tools. These results indicate that LI for tools and bodies are respectively negatively and positively correlated with the LI of faces, combining to produce the strongest relationship with face LI when directly contrasted. The slight increase in correlation magnitude for bodies*−*tools versus tools*−*bodies may be expected from the greater overlap in face and body responses (Figure S3). In contrast, tool selectivity is typically adjacent to face-selectivity and more likely to overlap body-selective responses, making direct parcel-to-parcel correlation a less perfect test of competition, since the rise of leftward laterality in parcel B may correspond more to a rise in rightward laterality in parcel A versus parcel B. Nevertheless, the neighboring and partially overlapping responses are associated with a negative correlation LI between faces and tools, and a positive correlation in LI between faces and bodies. The distributed pattern of correlations indicates that the co-fluctuation of laterality across these domains spans a large anatomical substrate. Interestingly, as shown in Figure S8, when we compared the faces*−*places contrast with tools*−*places contrast, we found that competition in VTC was masked by broad baseline-driven correlation; we were able to recover competition within the WM experiment only when additionally adding bodies to the baseline.

In summary, our analysis of the HCP data identified local lateralized competition between faces and tools, and local lateralized cooperation between faces and bodies, thereby supporting claims of the local-LNC theory.

## 3.5 Long-range coupling of laterality

We’ve provided evidence in line with certain forms of local competition and cooperation—competition between faces and tools, and cooperation between faces and bodies. However, the apparent relationship between reading onset and face laterality (Dundas et al., 2013, 2014; Behrmann and Plaut, 2020) was not explained in our data by local competition in VTC between representations of faces and words. However, beyond local constraint theories, other theories have emphasized the role of long-range connectivity in shaping the topographic and hemispheric organization of VTC (Rossion and Lochy, 2022; Price and Devlin, 2011; Mahon, 2022; Li et al., 2020). Here, we focused on the long-range-LNC theory (Rossion and Lochy, 2022), which states that the left lateralization of language has a primary long-range coupling effect with reading in VTC, and a primary local competition effect with social processing in a region of the posterior superior temporal sulcus (STSp) involved in language processing (Rajimehr et al., 2022); this local competitive effect with social processing is then argued to drive a secondary competitive effect of language lateralization on face lateralization in VTC, through a primary long-range coupling effect between VTC face representations and social processing in the STSp.

To test the predictions of this theory, we began by examining the coupling of laterality for text across a large distributed network activated by text, a network examined in our prior work with the same dataset (Vin et al., 2024).

Given that each node in this circuit exhibits significant leftward laterality (Vin et al., 2024), we predicted that the LI would also covary across the distributed circuit: that is, individuals with greater LI in VTC would also have greater LI in the other nodes of this circuit.

Following Vin et al. (2024), and as described further in Methods 2.1.5, we defined several anatomical regions of interest based on subsets of parcels from the Glasser atlas (Glasser et al., 2016); we used the same ROIs, with the exception of EVC, which we constructed as the union of V1, V2, and V3. Thus, we had six regions of interest: early visual (EVC), dorsal visual (IPS), ventral high-level visual (VTC), and language-related (IFG, IFGorb, PCG, STSG)^2^ (see Figure 7B and also Fedorenko et al., 2010, 2024). Within these regions, we computed selectivity and then LI. We then correlated the LI of each region with that of our VTC parcel. As shown in Figure 7C, we found that the LI in VTC is positively correlated with that of each region in this circuit (all *rho >* 0.33, all *p <* 0.05).

To ensure this pattern is not due to our particular choice of anatomical ROIs, we performed a finer-grained and more generic analysis using a symmetric atlas with 500 parcels in each hemisphere (Yan et al., 2023). We computed summed selectivity in each parcel, and then computed LI for all homotopic pairs of parcels (Figure 7D). Next, we computed the between-participant correlation of parcel LI with VTC laterality, using different subsets of runs (odd for VTC, even for other parcels). We plot these correlations, excluding VTC, in Figure 7E. To determine whether lateralized regions generally covary in their degree of laterality across participants, we computed the correlation between the average LI of each parcel, and its across-participant correlation with VTC. As shown in Figure 7F, we found a strong correlation between a parcel’s average LI and its correlation in LI with VTC across participants. Selecting a subset of the 50 most left-lateralized parcels, we found that these parcels have consistently high correlations in their LI with that of VTC (Figure 7G).

We performed a complementary analysis for faces in Figure S9. While we did find a significant correlation between the parcel LI and correlation with VTC LI (*rho* = *−*0.17*, p <* 0.001; the negative sign indicating that greater *rightward* parcel laterality corresponds to greater correlation in LI with VTC), this relationship is weaker than it is for words, and less visually apparent (Figures S9A,B,D). Rather than being driven by a large distributed circuit, as seen for text laterality, the coupling for faces appears to be driven mostly by a cluster of parcels in the posterior lateral temporal cortex, slightly posterior and superior to VTC and inferior to the STS.

In summary, these results provided support for a primary long-range coupling between frontotemporal language processing and word processing in VTC, which appears to percolate to other nodes of the word processing network including IPS and EVC. However, these results did not provide strong support for long-range coupling of face processing in VTC with other regions.

## 3.6 Long-range lateralized neural competition

Last, we assessed the claim of the long-range-LNC theory (Rossion and Lochy, 2022) that face lateralization emerges due to pressures to couple with downstream lateralized social processing regions, which are right-lateralized through competition with left-lateralized language processes (Rajimehr et al., 2022).

Given the broad coupling in laterality for text processing across frontotemporal areas, VTC, EVC, and IPS (Figure 7), a natural prediction of the long-range-LNC account is that selectivity for faces in VTC should compete with text laterality in these other regions. To assess this prediction, we adopted a data-driven approach similar to that used for the individual category whole-brain coupling analyses (Figures 7D-G, S9). In this case, however, we compared VTC face LI to the text LI of ROIs across the entire cortex, using the object baseline. We found little coupling in VTC face LI with broader cortical text LI (Figure S10). Moreover, we found no significant relationship between ROI-level text LI and VTC face LI, and the top text-lateralized ROIs show little consistency or strength in their correlation with VTC face LI.

Generally, these results do not support the long-range-LNC account, although it should be kept in mind that the perceptual demands of visual text processing—while activating downstream language regions (Vin et al., 2024; Stevens et al., 2017)—may not lead to robust engagement of linguistic processing mechanisms. Moreover, the shared object baseline may mask negative correlations, as discussed in the earlier section examining *reading-LNC*.

Two more direct predictions of the long-range-LNC account are that face LI should correlate negatively with language LI and positively with social LI (Rossion and Lochy, 2022). To test these predictions, we took advantage of the language HCP task, and a recently developed parcellation of language network regions used to demonstrate competition between social and language processing (Rajimehr et al., 2022), shown in Figure 8A and described further in 2.2.6. We then computed language and social LI within each ROI, using the "story" contrast for language, and the "theory of mind (TOM) *−* random" contrast for social processing, following (Rajimehr et al., 2022). Before proceeding to our analysis focused on VTC, we replicate the general results from (Rajimehr et al., 2022) that motivated the analysis, using our summation-based selectivity and the laterality index (L-R)/(L+R) rather than mean-selectivity and laterality difference (L-R) that were used in this prior work. First, shown in Figure 8B., we found broadly complementary group-level laterality indices for language in social processing across this network, with weaker LI magnitudes for social processing. Next, performing an inter-subject correlation of social and language LI in these regions, we replicated the finding of a negative correlation specifically in STSp, suggestive of a highly localized competition between social and language processing. Notably, areas 55b and PGi both show positive correlations, indicating that the relationship between language and social processing may not be exclusively competitive. Having replicated the social vs. language competition in STSp, we move on to test the predictions of the long-range-LNC account, by correlating social and language LI in each parcel with visual processing lateralities in VTC.

The results, shown in Figure 8C., demonstrate a significant negative correlation between language LI in PSL with both measures of face laterality, along with several trends towards negative correlations for other language network parcels. Additionally, we found that social processing laterality generally correlated positively with emotional face laterality in VTC, though these correlations did not survive family-wise statistical correction. Moreover, we found that VTC tool processing laterality was generally positively correlated with language laterality, particularly in STSa, while VTC body processing laterality followed a similar trend of negative correlation with language processing laterality in PSL, albeit not surviving family-wise statistical correction.

Noting that laterality is more variable among left-handers (Labache et al., 2023), we opted to take a broader look at the long-range laterality relationships across the full spectrum of handedness cutoffs, rather than just the right-handed population we have examined to this point. We repeated the same analysis as described above, at handedness thresholds spaced in scores of 10 from -100 to 90, where -100 corresponds to including all subjects, 90 corresponds to including only the most right-handed subjects, and 40 corresponds to the previous analysis. The results, plotted in Figure 8C, demonstrate a stronger negative correlation between language LI and VTC face LI at slightly weaker handedness thresholds, but otherwise, a minimal dependence on handedness threshold. Moreover, the relationships between VTC face LI and language LI in PSL survived after regressing out handedness, suggesting that they are not *explained by* handedness, but rather are dependent on having sufficient variability in lateralization, variability which is found across a more relaxed handedness cutoff (see also Vingerhoets et al., 2013). In contrast, the relationship between VTC tool LI and language LI generally increased when progressively including more left-handers. Assessing the relationships with social processing LI, we found that the negative correlation between STSp social LI and VTC emotional face laterality became significant at a slightly reduced handedness cutoff of 30, and remained similar for progressively relaxed cutoffs. A similar weak relationship was seen between VTC emotional face LI and STSa social LI at more relaxed handedness cutoffs. However, the handedness cutoff did not produce any significant correlations between the face *−* place VTC LI and social processing LIs. These results indicate a correspondence between face processing laterality in VTC and social processing laterality that is dependent upon the face processing task: a more socially-relevant emotional task was necessary to elicit this coupling in laterality. However, the results depended on a slightly weaker handedness threshold than our *a priori* choice.

In summary, the results provide weak but converging positive support for the long-range-LNC theory of face lateralization, through both negative correlations of face processing LI and language LI, and a positive correlation between specifically *emotional* face processing LI and social processing LI in the STSp, the same site where *local* competition is seen between social and language processing in the form of anti-correlated individual differences (Rajimehr et al., 2022).

### 4 Discussion

The current study examines individual variability in organization of visual categories within VTC, and leverages this variability to infer sources of its laterality. Although individuals show a generally similar arrangement in the global topographic organization of category selectivity in VTC (Kanwisher, 2010; Grill-Spector and Weiner, 2014), we demonstrated substantial individual variability in fine-grained category-selective topographic organization and hemispheric laterality, which are stable across individual measurements within individuals but much more variable across individuals. These findings suggest that various forces may cause differences in organization across development (Behrmann and Plaut, 2020). In the current work, we tested the predictions of developmental theories which propose *lateralized neuronal competition* (LNC), relying on individual differences to infer the mechanisms of functional interactions, and examining both local and long-range competitive/cooperative factors.

## 4.1 No evidence for graded lateralized competition between words and faces in VTC

First, we tested the prediction of Neuronal Recycling (NR) (Dehaene and Cohen, 2007) and Graded Hemispheric Specialization (GHS) (Behrmann and Plaut, 2020)—two theories captured under the umbrella of reading-related lateralized neural competition (reading-LNC) theories (Rossion and Lochy, 2022)—of a competitive relationship in lateralization for written text and faces in VTC. While it is known that both face and text selectivity can have flipped dominance in the case of rightward language laterality (Gerrits et al., 2019)—which is more common, albeit still rare, in left-handed individuals—here we sought to test competition within right-handed individuals with gradations in the left lateralization of text selectivity. Across three different contrasts and three different selectivity measures in a relatively large group of proficient readers, we found no evidence of a competitive relationship between faces and text. Rather, if anything, we found what seems to be a weak positive correlation between the LI of faces and text, and a stronger positive correlation between the LI of faces and objects.

The lack of apparent competition in lateralization for any of the categories is surprising (Dehaene et al., 2010; Dundas et al., 2013, 2014, 2015; Centanni et al., 2018), but replicates another fMRI study (Davies-Thompson et al., 2016) that computed LI based on either peak response or number of selective voxels. Our larger sample, assessment of more contrasts, and assessment of the additional summed selectivity measure—which accounts for both the strength and size of the selective region—thus strongly corroborates this prior study (Davies-Thompson et al., 2016). While our results appear to be directly at odds with the study of Centanni et al. (2018), which found a negative correlation in the size of left VWFA and left FFA in children learning to read, this study used confounding baselines, comparing the size of regions defined with opposite contrasts (letters > faces for VWFA, faces > letters for FFA). Thus, it is entirely possible that the category-based confound explains their effect, rather than a true competition between letter and face responses. Moreover, our results are consistent with a recent study that found no correlation in the LI of face and word selectivity in either children or adults (Liu et al., 2024). Last, our results are also in line with recent longitudinal studies demonstrating that emerging text representations do not recycle face-selective cortex, but rather appear to emerge in previously non-selective cortex (Dehaene-Lambertz et al., 2018) or weak, waning limb-selective cortex (Nordt et al., 2021; Kubota et al., 2023).

## 4.2 Evidence for more general local interactions

Given the lack of evidence for local competition between faces and text, we wondered whether the laterality of faces might instead be related to individual laterality patterns across a range of other categories in VTC. Notably, individual variability in topographic organization for faces and other categories in VTC can be predicted with models that consider shared information spaces with individual spatial tuning, in a way that generalizes to new stimuli (Haxby et al., 2020; Feilong et al., 2023), in line with distributed and overlapping information patterns in VTC (Ishai et al., 1999; Haxby et al., 2001). Thus, local cooperation and competition between representations involved in the perception of a much broader range of categories may influence face laterality on an individual basis; we termed this idea the local-LNC theory, which, while emphasizing generic local representational competition, is not mutually exclusive with the possible role of reading (reading-LNC) or long-range coupling (long-range-LNC) in shaping VTC laterality.

We first found a strong correlation in the LI for face and object selectivity in VTC, across the held-out category baseline, the fixation baseline, and the fixation baseline after regressing the held-out category vs. fixation LI; this correlation was not seen, however, when using the full baseline, including a category-based negative confound (i.e. faces - objects + text vs. objects - faces + text). A weaker correlation was also seen between faces and text that survived the same 3/4 contrasts, however it was about half the magnitude of that between faces and objects. In contrast, a positive LI correlation between objects and text was present only for the raw fixation baseline, implying it is not a true correlation. The strong LI correlation between faces and objects is in line with the local-LNC theory, and in particular with the finding of a stronger cost in simultaneously represented faces and objects, compared to other more neurally distinct categories (e.g. faces and scenes), indicating representational overlap (Cohen et al., 2014).

Next, an analysis of data from the Human Connectome Project revealed distinct pairwise relationships in LI among visual categories, with face LI positively correlated with body LI and negatively correlated with tool LI, even across independent tasks. Local (anti-)correlations were widespread in VTC, and notably strong within the fusiform face complex (FFC) anatomical parcel, the most frequent site of face selectivity. When we constructed explicit contrasts of bodies versus tools, the magnitude of relationships with face LI was enhanced relative to either individual category contrasted against places; moreover, the relationship was stronger for LI of bodies versus tools than for LI of tools versus bodies, in line with the greater overlap in selectivity for faces and bodies.

What might explain the relationships between faces, bodies, and tools? Notably, body selectivity overlaps strongly with both face selectivity (Downing et al., 2006; Peelen et al., 2009) and tool selectivity (Bracci et al., 2012), which have a complementary lateralization profile. In particular, tool and *hand* selectivity overlap in the lateral fusiform gyrus (Bracci et al., 2012). That the viewing of faces frequently co-occurs with viewing bodies, and both inform person recognition and social processing, may give rise to the partially overlapping responses in VTC. Similarly, the co-occurence of viewing hands and tools—both one’s own, and those of others observed using tools—may give rise to the partial overlap in their functional organization. While hand—and more broadly *limb* (Weiner and Grill-Spector, 2010)—selectivity overlaps with body selectivity, body selectivity overlaps more strongly with face selectivity (Peelen et al., 2009), compared to the weaker overlap of limb and face selectivity (Weiner and Grill-Spector, 2010); however, in HCP, the bodies condition contains limbs, hands, and headless bodies, so finer-grained conclusions cannot be drawn. Moreover, prior studies make clear that the functional organization of VTC is graded, with each category-selective area falling in a particular preference zone that partially explains responses to multiple categories (Konkle and Caramazza, 2013; Bao et al., 2020; Yao et al., 2023). The local-LNC account claims that this graded topography is the product of cooperative and competitive pressures occurring locally, albeit influenced by both bottom-up and top-down demands. This idea is in line with computational models that consider VTC as a unitary representational space with graded topographic specialization encompassing selectivity for (and representations of) both categories and other visual features, which evolve in an inherently dependent fashion (Blauch et al., 2022b; Doshi and Konkle, 2023), even if the outcome is a large degree of specialization (Blauch et al., 2022b; Dobs et al., 2021).

One challenge to the local-LNC theory comes from the study of (Cohen et al., 2014). Whereas their finding of a stronger cost in simultaneously representing faces and objects—driven by greater neural representational overlap—is in line with our results of correlated LI between faces and objects, they also found a weaker cost in simultaneously representing faces and bodies, which appears at odds with our results of correlated LI between faces and bodies. Similarly, the face/body distinction has been proposed as a strong organizational principle for VTC (Yargholi and Op de Beeck, 2023). Nevertheless, face and body representations have been shown in several studies to strongly overlap (Downing et al., 2006; Peelen et al., 2009), and both score high on the animacy dimension, which is proposed as one of the two main organizing dimensions of VTC (Yargholi and Op de Beeck, 2023; Konkle and Caramazza, 2013; Bao et al., 2020). Notably, self-supervised deep neural networks that learn visual representations without semantic tasks show a particular relationship between face and body responses; specifically, lesioning face-selective units leads to a strong cost in representing the categories that exhibit the strongest cost when lesioning body-selective units (Prince et al., 2024). This implies that the visual statistics of viewing faces and bodies is sufficient to drive meaningful overlap in their neural responses, which appears to manifest as shared laterality. However, the body stimuli in the study of (Cohen et al., 2014) can be distinguished largely based on the limbs, which have a more separable neural representation with faces (Weiner and Grill-Spector, 2010), perhaps explaining their finding of a weaker cost in simultaneous representation of faces and bodies.

## 4.3 Long-range coupling of laterality

Last, we explored long-range coupling as a mechanism for the emergence of VTC lateralization. It is widely accepted that the lateralization of language drives the lateralization of text processing in VTC (Dehaene et al., 2010; Li et al., 2020). Accordingly, VWFA hemispheric dominance has been shown to track the hemispheric dominance of language in left-handers with greater variability in language dominance (Gerrits et al., 2019). However, whether variability in VTC text lateralization similarly corresponds with that in language areas has not been explored within the more modest variability seen among right-handers. Here, we found that a large distributed and lateralized cortical circuit of regions involved in text processing (Vin et al., 2024) showed correlated text laterality with VTC across subjects. The correspondence in laterality between VTC and frontotemporal regions corresponds well with prior work using functional and structural connectivity with language areas to predict the location of the VWFA (Saygin et al., 2016; Stevens et al., 2017; Li et al., 2020). However, the correspondence with IPS and EVC is more surprising (but see Kay and Yeatman, 2017), suggesting that text processing engages a highly recurrent and interactive cortical circuit in which language lateralization percolates to text processing in all of the nodes. Whether the lateralization of processing in EVC and IPS emerges temporally before or after top-down feedback is an interesting question that could be addressed with more time-resolved methods such as MEG; if in advance of top-down feedback, it would suggest that the bottom-up tuning of these regions has been shaped by the lateralization of language, as is thought to be the case for text selectivity in VTC (Dehaene and Cohen, 2007; Price and Devlin, 2011).

Given this finding of long-range coupling in laterality for text, we next sought to explore the direct predictions of the *long-range-LNC* account regarding face lateralization: that is, face LI should negatively correlate with language LI, and positively correlate with social processing LI in homotopic frontotemporal areas encompassing the typically left-dominant language network (Rossion and Lochy, 2022). Indeed, broadly complementary laterality patterns for social and language processing have recently been found across the language network, with a negative correlation specifically in the STSp (Rajimehr et al., 2022); our goal was to test whether this may explain variation in the laterality of face processing in VTC.

We performed a targeted analysis of the same language network parcels explored by (Rajimehr et al., 2022), finding that VTC LI for both face processing tasks was negatively correlated with language processing LI in the PSL, and positively correlated with social processing LI in STSp, only for the emotional face processing task. These results are broadly in line with the predictions of the *long-range-LNC* account. However, in both cases, the effects were somewhat weak and sometimes dependent on using a slightly more relaxed handedness cutoff (although not requiring the inclusion of left handers). These weak effects would be expected to be quite difficult to discover in smaller studies, which might partially explain our inability to find a correlation in non-VTC text LI with VTC face LI in our in-house data. However, in the case of social processing, the small effect sizes are likely also due to the imperfect match between the social demands required in HCP task (simple theory-of-mind judgments regarding the motion of simple shapes), and the social demands required in typical real-world face processing scenarios. Similarly, the language coupling results might be weakened by the use of an auditory language localizer task, rather than a visually-based one. While we are not aware of any studies that have directly compared language network LI in corresponding visual and auditory tasks within individuals, both forms of the task have been used across studies of the language network (Fedorenko et al., 2024). To the extent that differences exist across tasks, however, it might be expected that the visually-induced laterality would couple more strongly with visual lateralities in VTC.

In exploratory analyses, we found somewhat stronger relationships between language LI and the LI for tools, as compared to faces. In line with our findings, tool processing has generally been found to be left lateralized in humans (Chao et al., 1999; Downing et al., 2006; Johnson-Frey, 2004; Lewis, 2006), reflecting personal handedness and use but also the statistics of the handedness of viewed tool use (Gainotti, 2015). It also corroborates previous findings that distributed LI patterns for tool pantomiming and verb generation are highly correlated across individuals with typical and atypical language lateralization (Vingerhoets et al., 2013). Indeed, the left lateralization of neural mechanisms underlying tool processing has been proposed as a causal link in the evolution of a left-lateralized speech controller (Frost, 1980; Stout and Chaminade, 2012). Additionally, in our exploratory analyses we found a trending correlation between VTC body LI and PSL language laterality, similar to the relationship between PSL and VTC face LI; however, no relationships were seen between body LI and social LIs.

## 4.4 Limitations

Our results suggest that there is local lateralized competition between faces and tools in VTC, but not between faces and text, and long-range coupling between faces and social processing, as well as between text and tools and language processing. However, there are some important limitations that warrant caution in an overly strong interpretation of our results.

First, with respect to our null findings for reading-LNC predictions, responses for objects, faces, and text appear to meaningfully overlap—not just in estimated responses (which could be due to overlapping hemodynamic responses), but also in selectivity relative to a scrambled image baseline. This representational overlap made it difficult to construct unbiased selectivity contrasts. In general, given three categories *A, B, C*, when comparing the LI of category *A* and *B*, there are four natural options: 1) compare *A* and *B* directly, 2) compare *A−C* and *B−C*, using a shared baseline to account for general activation-increasing effects such as arousal, 3) compare *A−B* and *B−A* 4) compare *A−(B+C)* and *B−(A+C)*, using the “full” baseline which would make up a standard category selectivity contrast.

None of these choices is ideal: option 1 yields strongly positive correlations, due in part to the imperfect ability of the GLM to fully identify the true sources of activation, and thus partially blurring responses across different categories; option 2 induces a bias towards positive correlation due to the shared baseline; option 3 may be meaningful due to the asymmetry imposed by summation of positive selectivity, but induces an obvious strong bias towards negative correlation; and option 4 exhibits competing biases towards negative and positive correlation, which do not simply magically cancel out.

By using independent splits of data to compute each contrast, we were able to remove statistical bias in our correlations of LI, but not category-based biases arising from shared categories across compared selectivity contrasts. We used options 1, 2, and 4 in our analyses of faces, text, and objects (Figure 4. None of the options revealed competition between faces and text, whereas only option 4 revealed competition between text and objects; however, this competition between text and objects appeared to be driven by the shared (face) baseline, as it was eliminated when either option 1 or 2 was used. Competition was not found when we replaced the sum selectivity metric with either of two more standard options: peak voxel selectivity, or number of selective voxels.

In line with these ideas, the demonstration of anti-correlated LI between tools and faces relied on having entirely independent contrasts (e.g., no overlapping conditions) across two separate experiments (WM and Emotion). When we used overlapping contrasts in HCP (faces*−*places vs. tools*−*places), the competition was not observed (however, in this case, statistical dependence could not be eliminated, as contrasts were built across only two runs collected in opposite phase-encoding directions). Additionally, the substantially larger size of HCP provides greater statistical power to discover the competitive relationships of interest, so the results from the two experiments cannot be compared directly. That said, the lack of trending competitive effects between face and text selectivity in the main experiment, and the corroboration of our results with prior studies (Davies-Thompson et al., 2016; Nordt et al., 2021), suggests that statistical power was likely not an issue.

Last, while there is a clear connection between our results and prior literature on the overlap in mechanisms for tool and language processing, the associations between tool and language LI in this experiment must be viewed with caution. Many of the tool images contained tools with textual brand labels (e.g., DeWalt), implicitly encouraging participants to verbally recode while viewing some of the tool images and thereby driving language processes in the brain. To the extent that such language processes activate VTC (e.g. Woolnough et al., 2021; Li et al., 2023), the linguistic—rather than tool-based—properties of tool images might drive both the correlation in LI with language areas, and the anti-correlation in local LI with face processing.

## 4.5 Combining ideas from different theoretical accounts

Why might faces exhibit local competition with tools but not text? One relevant factor is the relatively late acquisition of reading, which occurs after face processing and selectivity has already substantially developed (Nordt et al., 2021; Kosakowski et al., 2022), albeit certainly not fully (Scherf et al., 2007; Peelen et al., 2009). In contrast, humans explore manipulable objects and have significant experience viewing body parts and most other broad visual categories at a young age; indeed, words are an exceptional visual category insofar as visual expertise is introduced at a relatively late point in life. Prior work has demonstrated that the acquisition of the Devenagari script introduces new text response patterns without strongly interfering with pre-existing response patterns for other categories (Hervais-Adelman et al., 2019), a finding deemed “recycling without destruction”. This is in line with the overlap we found in VTC category selectivity versus a scrambled baseline (Figure S1), suggesting that faces and text may not compete in the way that has been previously suggested (Behrmann and Plaut, 2020), due to greater overlap than commonly appreciated. Indeed, a recent intracranial study in humans found that 39 % of word-selective electrodes in the right hemisphere and 47 % of such electrodes in the left hemisphere could discriminate faces from the other object categories excluding words (Boring et al., 2021), indicating overlap in representations at a spatially precise level. Given this overlap, it would be interesting to re-examine how the emergence of word *responses*, rather than word *selectivity*, relates to prior face selectivity, during the acquisition of reading Roman script in English and French children in the datasets collected by (Nordt et al., 2021) and (Dehaene-Lambertz et al., 2018), respectively. Examining only the *word-selective* voxels may inherently exclude those voxels which gain new responses to words but retain strong responses to faces (Dehaene-Lambertz et al., 2018; Nordt et al., 2021).

If words do not induce local competition with faces in VTC, the previous findings of a link between literacy and face lateralization (Dehaene et al., 2010; Dundas et al., 2013, 2014, 2015) need to be explained in other terms. The language-related or long-range-LNC account (Rossion and Lochy, 2022) was proposed to do just this; however, it does not immediately explain why face lateralization increases developmentally after the onset of literacy. One relevant factor is the broad cortical changes that are seen when learning to read, including in VTC, but also in occipital, superior temporal, and frontal areas (Dehaene et al., 2010; Monzalvo and Dehaene-Lambertz, 2013; Chyl et al., 2018; Hervais-Adelman et al., 2019). Additionally, the acquisition of reading literacy is usually accompanied with the acquisition of writing, which may compound the effects on distributed brain circuits (Hervais-Adelman et al., 2022). These global changes might enhance distributed competitive lateralizations between text-related (e.g., language) and face-related (e.g., social) processes, giving rise to an increase in long-range pressures on face laterality with the emergence of reading, even without local recycling or competition. This idea is supported by a concurrent study to ours, which demonstrated a negative correlation between sentence reading LI in Broca’s area (i.e. IFG) and face LI in VTC in adults but not children, where the LI in Broca’s area also increased over development (Liu et al., 2024).

We must emphasize that the finding of local (anti-)correlations in laterality indices between faces and tools/bodies are not inconsistent with the idea that these patterns are driven by long-range coupling. Indeed, as shown in Figure 1C, within the local-LNC theory, some long-range force is necessary to instigate a primary lateralization which would then influence secondary lateralizations through local interactions. The key difference between long-range and local LNC is that whereas long-range LNC proposes that the competition occurs locally at the distal site (e.g. STSp, between language and social processing), local LNC proposes that both competition and cooperation occur locally due to viewing statistics and representational characteristics. Of interest, one study demonstrated that both face and body representations in higher-level visual cortex show reduced rightward lateralization in left-handers (Willems et al., 2010), in line with a greater prevalence of atypical rightward language lateralization in left-handers and associated reductions in tool laterality in left-handers (Vingerhoets et al., 2013).

Based on our findings and the prior works discussed here, we suggest that local and long-range factors work synergistically to produce individual laterality patterns, as illustrated in Figure 9. As suggested by long-range LNC and reading-LNC, VTC word laterality is inherited through long-range coupling with left-lateralized frontotemporal language mechanisms. As suggested by long-range LNC, local competition between language and social processing happens primarily in the STSp, leading to rightward lateralization of social processing, and the emergence of rightward VTC face laterality due to long-range coupling with social processing in the STSp. However, as suggested by local-LNC, the results of these long-range coupling influences are constrained by overlap and interactions of local representations across a broad swath of VTC. While both the long-range and local interactions remain incompletely specified, we believe attention to both of them is key to fully understand the mechanisms of lateralization in VTC. Future work will be required to untangle the more precise contributions of both local and long-range factors.

**Figure 9:**
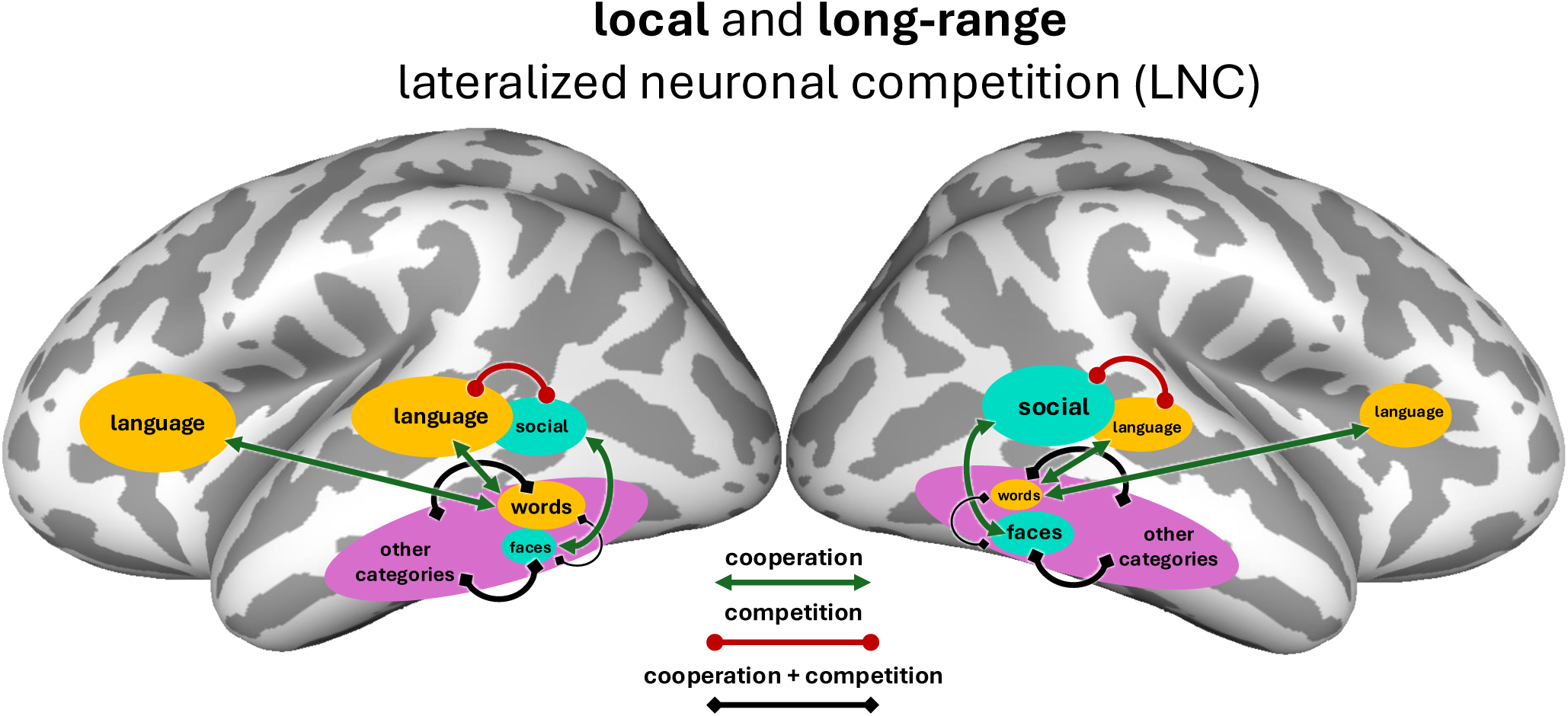
Schematic for the local and long-range lateralized neuronal competition theory, incorporating facets of both the local and long-range LNC accounts, based on the results of this study. The schematic emphasizes 1) that long-range coupling between left-lateralized language (in frontal and temporal areas) and words (in VTC) contributes to the lateralization of words in VTC, 2) that exclusive competition between words and faces is unlikely to be a major driver of rightward face laterality; however, a mix of competition and cooperation between words and faces is expected due to their partially overlapping responses, as shown with a thin black line, 3) social processing, which may become right-lateralized in the right STSp due to local competition with language induces a rightward long-range lateralization pressure on face representations, 4) words and faces both exhibit a mix of local competition and cooperation with neighboring visual representations in VTC. This theory schematic is not meant as an exhaustive list of all of the interactions shaping the lateralization of VTC, nor all of the areas involved in processing the various stimuli and tasks; rather, it highlights that competition and cooperation at both local and long-range scales influence the laterality of VTC.

## 4.6 Conclusion

Our work explores the principles of local representational competition and cooperation and long-range coupling with domain-relevant processing systems—principles which are essential to the theories of graded hemispheric specialization (Behrmann and Plaut, 2015, 2020; Plaut and Behrmann, 2011) and neuronal recycling (Dehaene and Cohen, 2007; Dehaene et al., 2010). However, it calls into question the claim made by both of these "reading-LNC" theories that VTC face laterality emerges specifically as a result of local competition with words, joining recent works (Nordt et al., 2021; Rossion and Lochy, 2022; Kubota et al., 2023). While our results do not strictly rule out this competition, we find evidence only for local competition with tools, vis-a-vis the negative correlation between tool and face LIs within VTC. Instead, VTC face LI generally appears to be correlated with VTC object LI, body LI, and even text LI in some analysis approaches. Assessing the possible role of coupling in lateralization with distant regions, we find evidence that VTC face LI is both negatively correlated with language LI in PSL, and positively correlated with social processing LI in the STSp. Our work is thus consistent with both local and long-range lateralized neuronal competition. We speculate that laterality patterns between other categories are similarly coupled locally and with downstream areas—even when those categories show weak group-level LI—with gradations across categories related to the overlap in their input and output demands. Future work may be able to take advantage of existing datasets to test this idea using the methods developed here, specifically focused on the full profile (size and strength) of selectivity. Additionally, the success of large-scale naturalistic imaging paradigms in elucidating the nature of visual representations in human VTC (Allen et al., 2022; Chang et al., 2019) motivates the use of similar approaches in a larger cohort of subjects to facilitate individual difference analyses that could provide greater clarity on the coordinated development of topographic and hemispheric organization, with particular relevance to the more generic or feature-based competition inherent to ideas about local interactions. Last, advancing computational models with increasingly realistic visual inputs (Russakovsky et al., 2015; Gan et al., 2020), processing mechanisms (Yamins et al., 2014; Kubilius et al., 2018; Spoerer et al., 2017), tasks (Bakhtiari et al., 2021; Mineault et al., 2021; Nayebi et al., 2023; Zhuang et al., 2020; Konkle and Alvarez, 2022), and spatial embedding (Blauch et al., 2022b; Keller and Welling, 2021; Achterberg et al., 2022; Margalit et al., 2024), will make it increasingly possible to make quantitative predictions about representational cooperation and competition *in silico*, which may reduce demands on—and aid the interpretation of—costly future experiments.

## Acknowledgments

We thank Marge Maallo for help with pilot funding acquisition, data collection and for useful early discussions. Additionally, we thank past and present members of the Behrmann Lab, along with Leila Wehbe and Michael Arcaro, for useful feedback throughout the course of the project. Last, we thank Kelly Martin and Jacob Prince for feedback on the manuscript.

## Data and Code Availability

Upon publication, all code and data will be made publicly available. Data will be made available on KiltHub, and code will be made available on GitHub.

## Author Contributions

**Nicholas M. Blauch**: Conceptualization, Methodology, Software, Validation, Formal analysis, Investigation, Resources, Data Curation, Writing - Original Draft, Writing - Review & Editing, Visualization, Supervision, Project administration, Funding acquisition. **David C. Plaut**: Conceptualization, Methodology, Writing - Review & Editing, Supervision, Funding acquisition. **Raina Vin**: Conceptualization, Methodology, Software, Formal analysis, Writing - Review & Editing. **Marlene Behrmann**: Conceptualization, Methodology, Writing - Review & Editing, Supervision, Project administration, Funding acquisition.

## Footnotes

1 Due to the smaller available sample with a scrambled condition (n=27), we do not use these maps for studying individual differences.

2 While we did not perform functional localization of language-selective voxels within these areas, our use of summed selectivity sidesteps the issue of aggregating unrelated neighboring responses, since negative preferences in the same parcel do not influence this measure.

