## Supplementary Information for "Individual variation in the functional lateralization of human ventral temporal cortex: Local competition and long-range coupling"


#### 1 Group category-selective maps

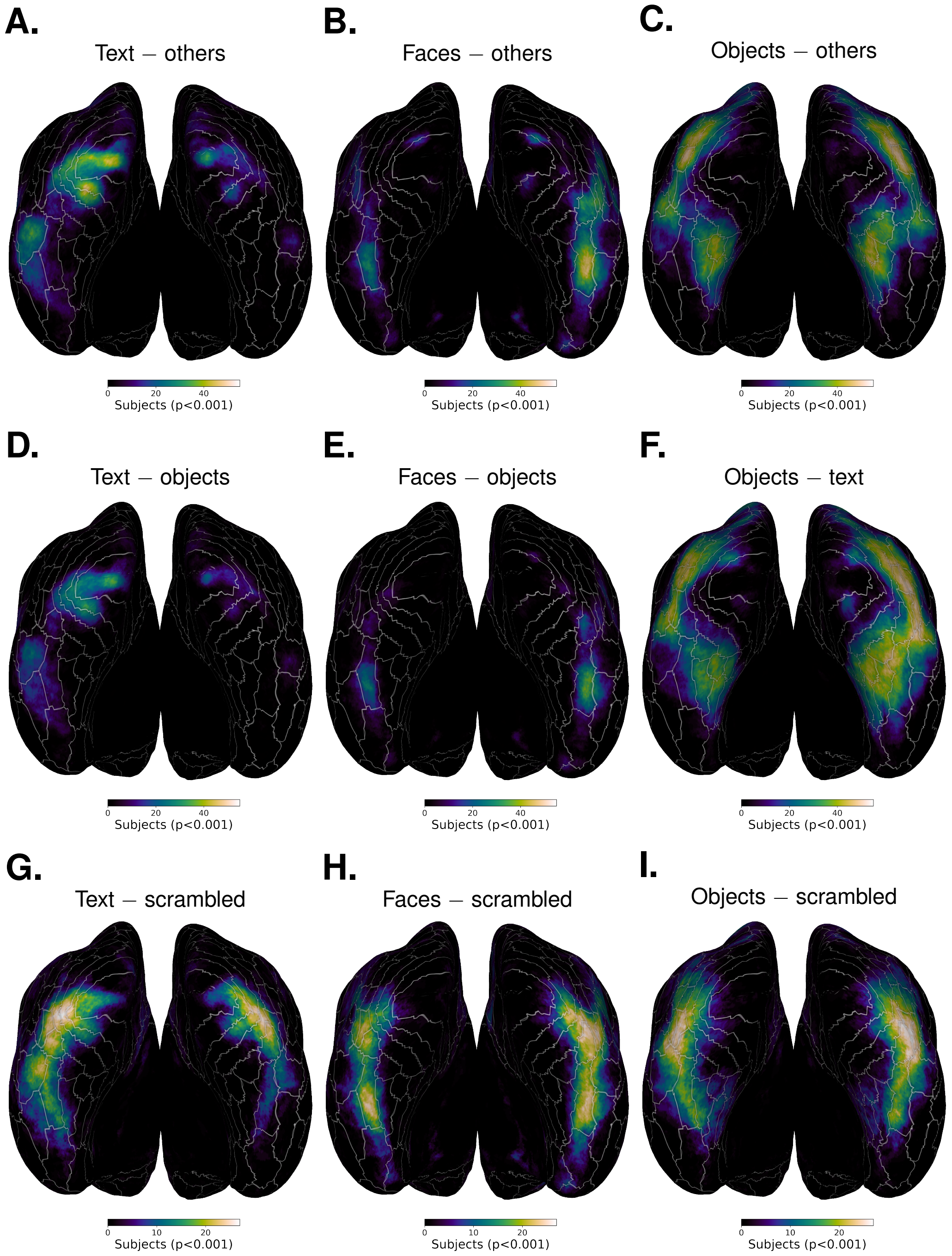


**Figure S1:** Group selectivity maps for relevant contrasts in the main experiment. Each map shows the number of participants that yielded significant selectivity for a given vertex in group surface space, where significance was determined as the selectivity value having $p<0.001$. Note: 55 participants were scanned, but only 28 saw the scrambled condition used in the second row.

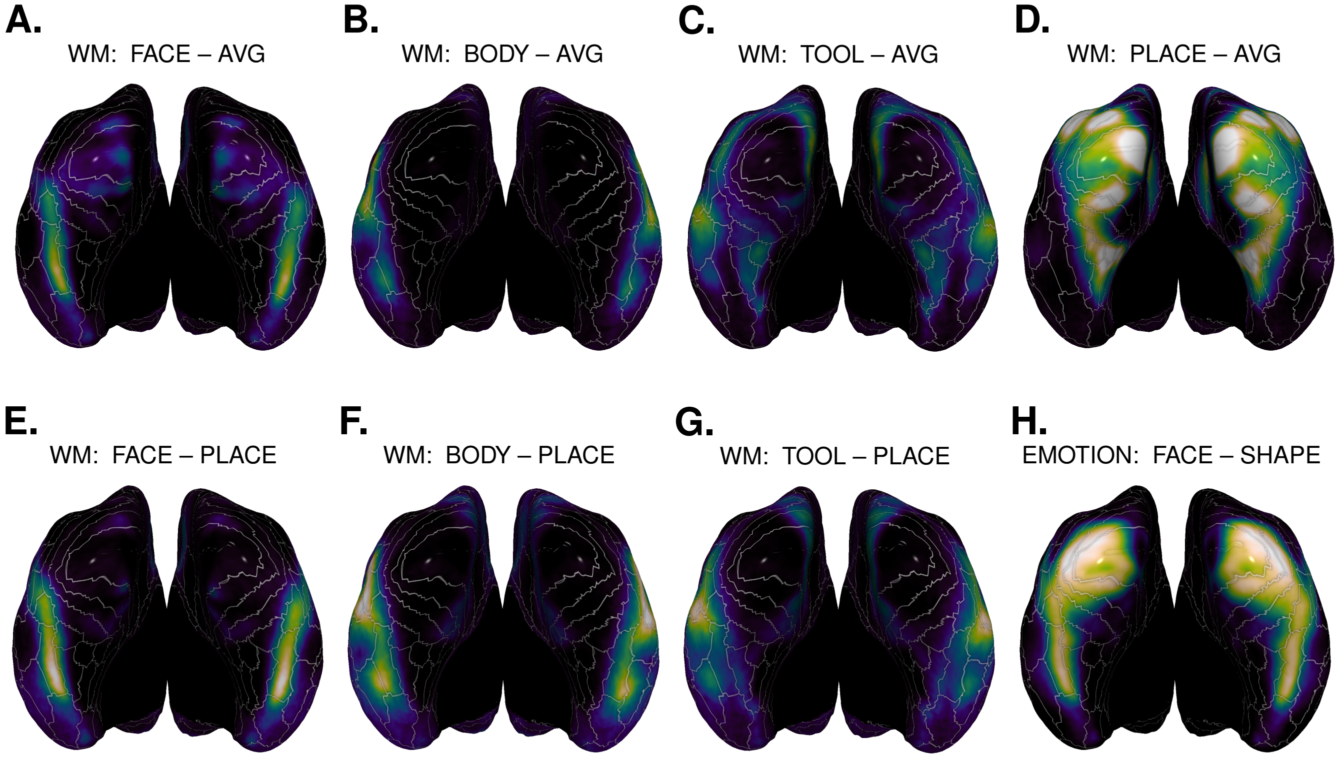


**Figure S2:** Group selectivity histograms for HCP contrasts. Color map goes from 0 to 700 participants showing selectivity ($p < 0.001$).

#### 2 Overlap of category-selective responses

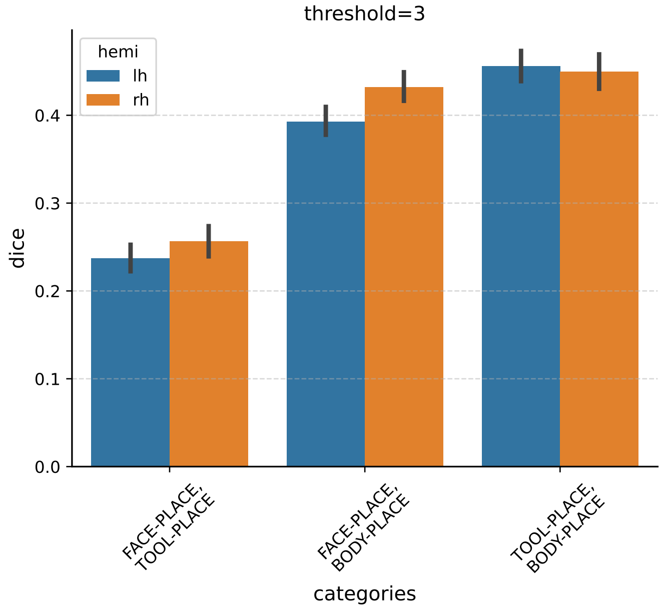


**Figure S3:** Overlap of category-selective regions in VTC, using the HCP WM contrasts of faces, tools, and bodies against places. A threshold of 3 was used to binarize the contrast maps before computing the dice coefficient.

### 3 Laterality using different metrics of selectivity

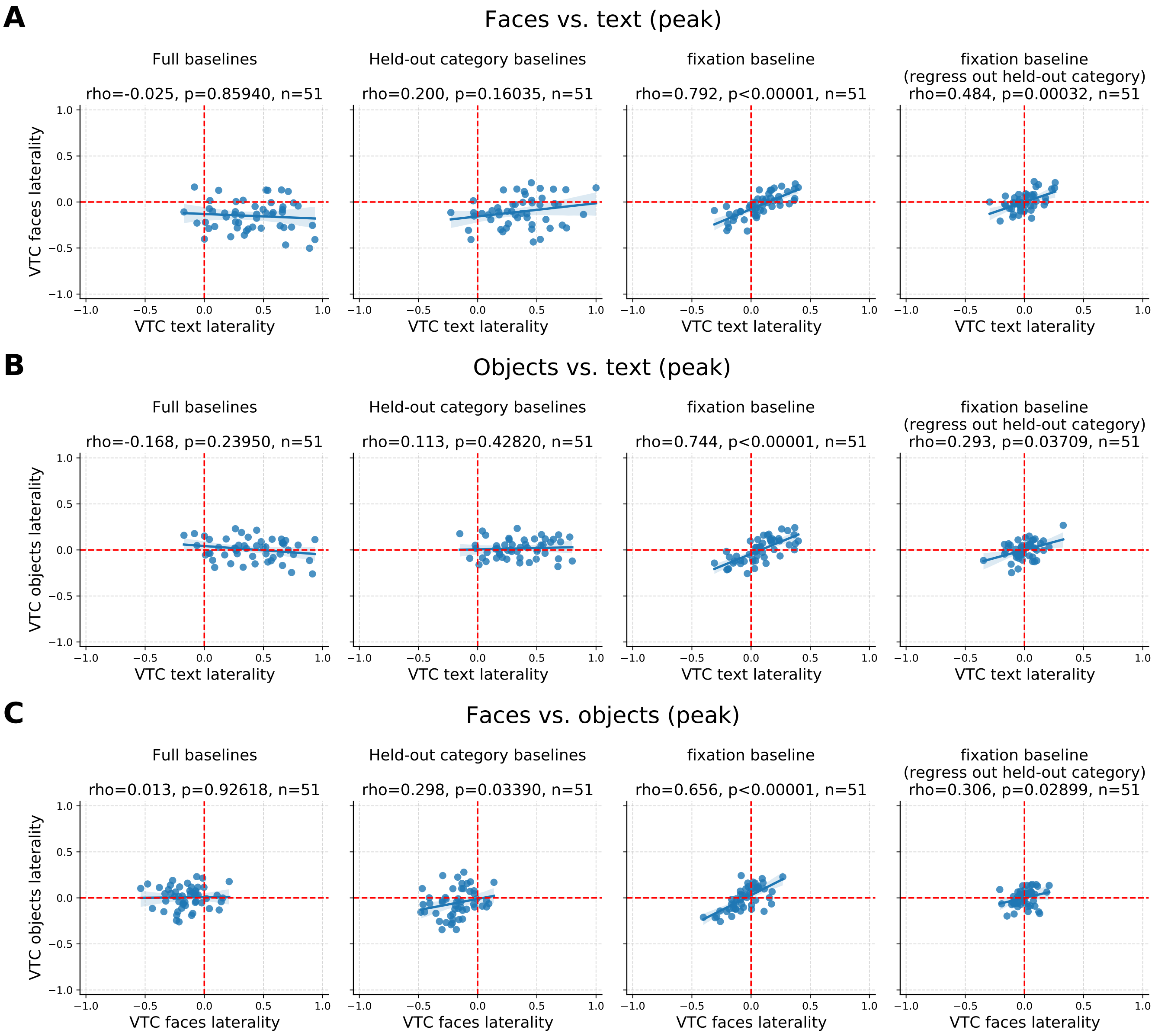


**Figure S4:** LI correlations using peak selectivity

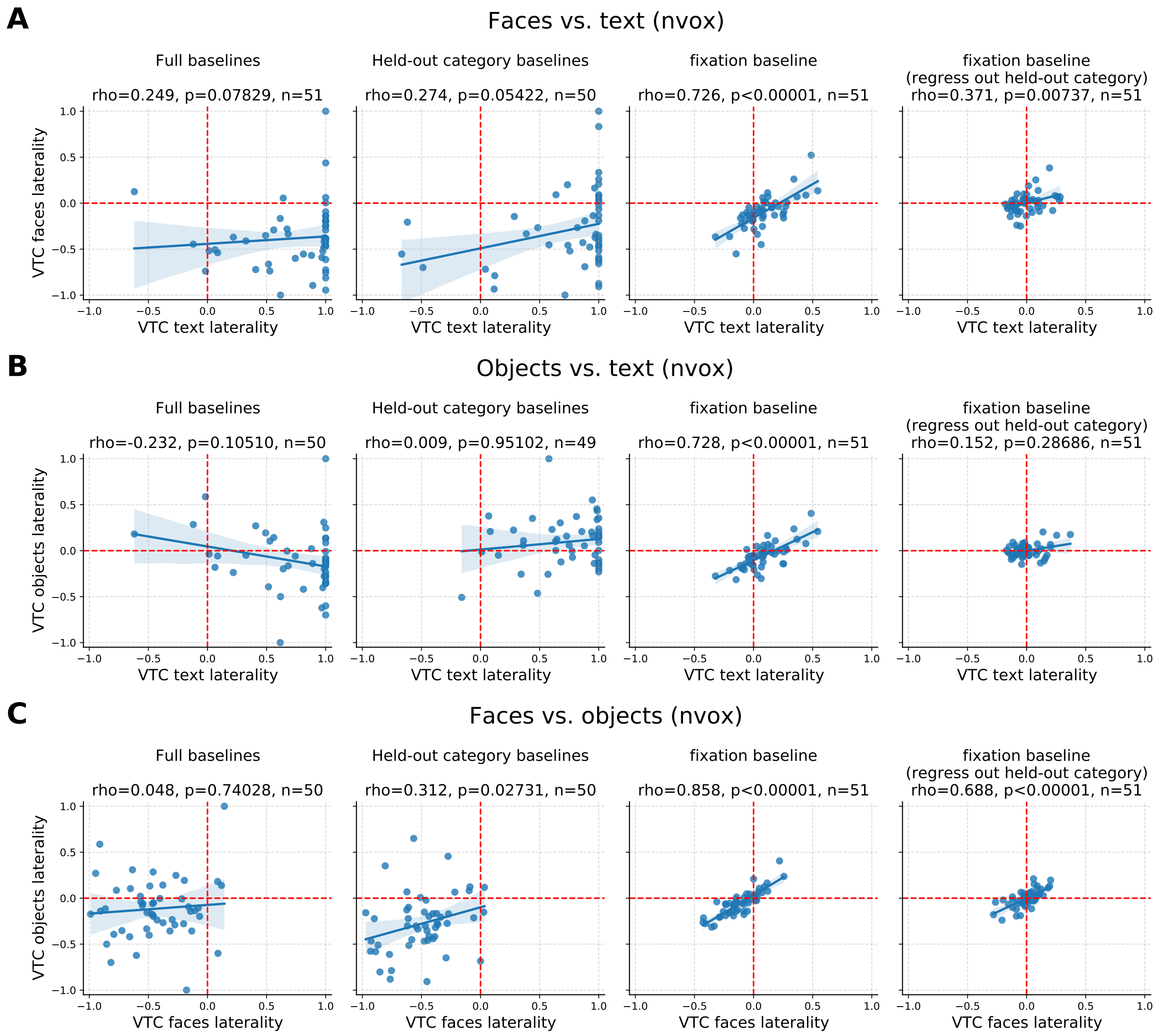


**Figure S5:** LI correlations using the number of selective voxels as a selectivity metric

### 4 Individual category-selective maps

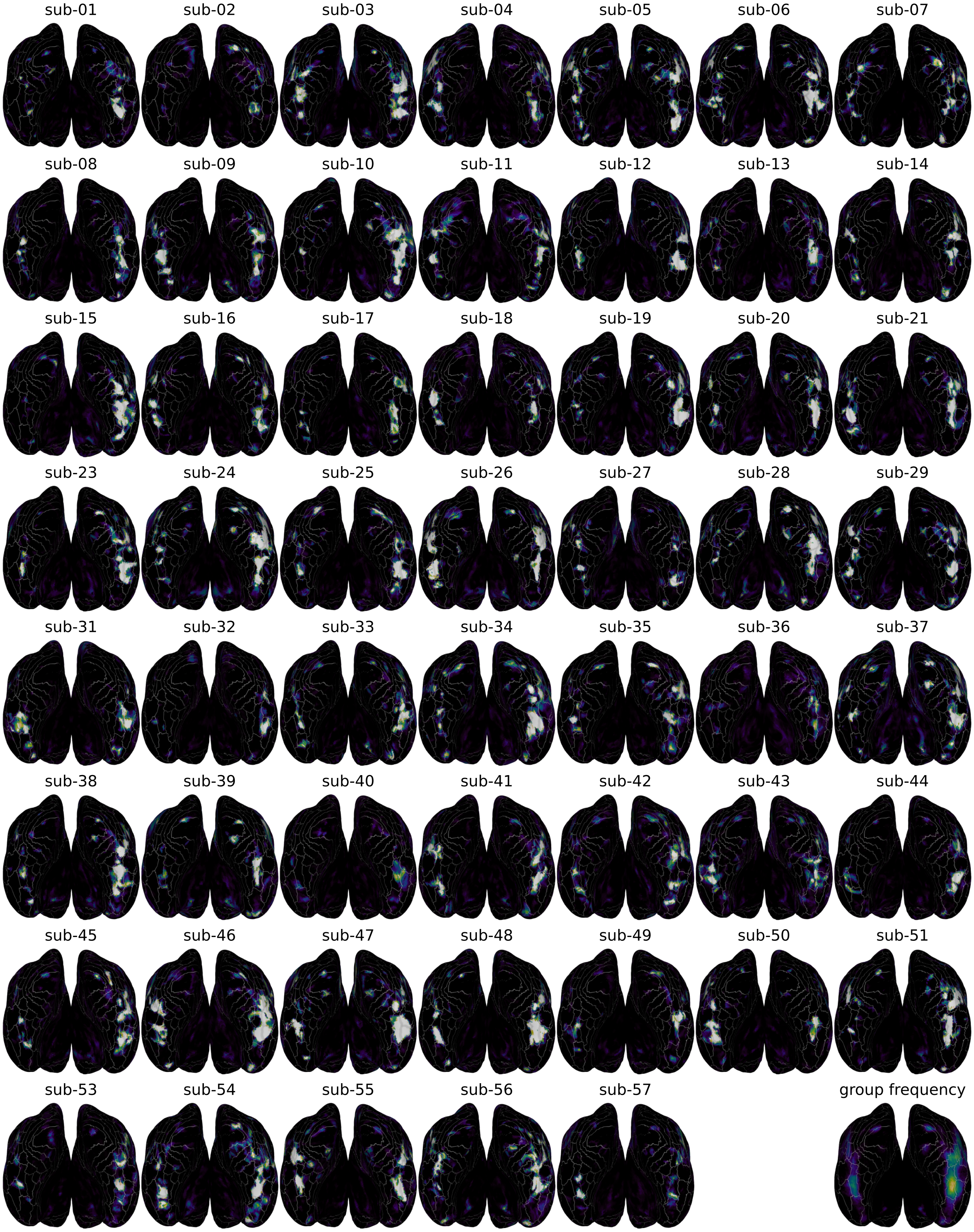


**Figure S6:** Individual face-selective maps for all subjects in our in-house experiment. The same approach as used in the main text (Figure 4D). was used to generate these maps, except all 5 runs (rather than even or odd) were used.

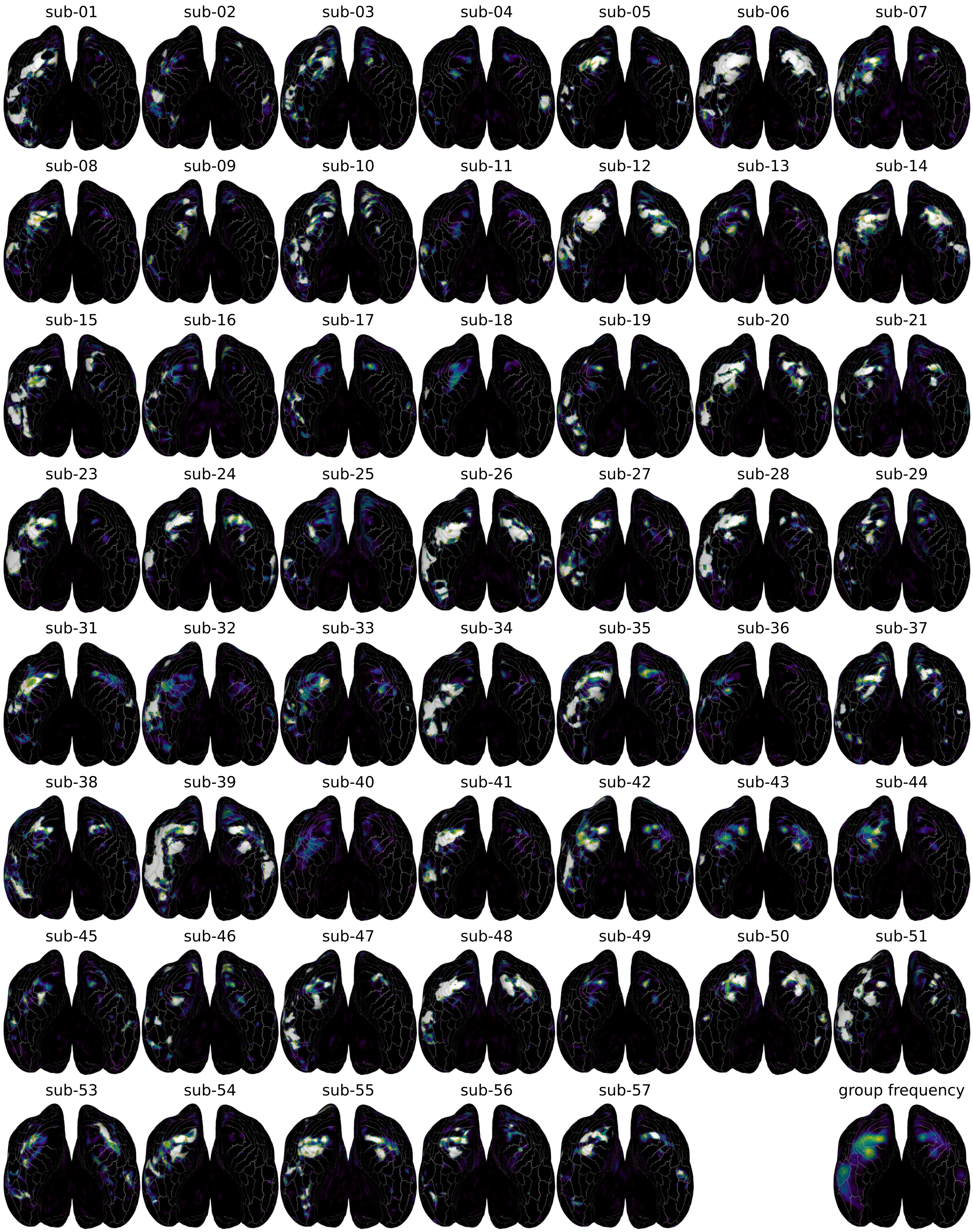


**Figure S7:** Individual text-selective maps for all subjects in our in-house experiment. The same approach as used in the main text (Figure 4D), except all 5 runs (rather than even or odd) were used.

#### 5 Additional contrasts for parcel-level LI correlations in HCP

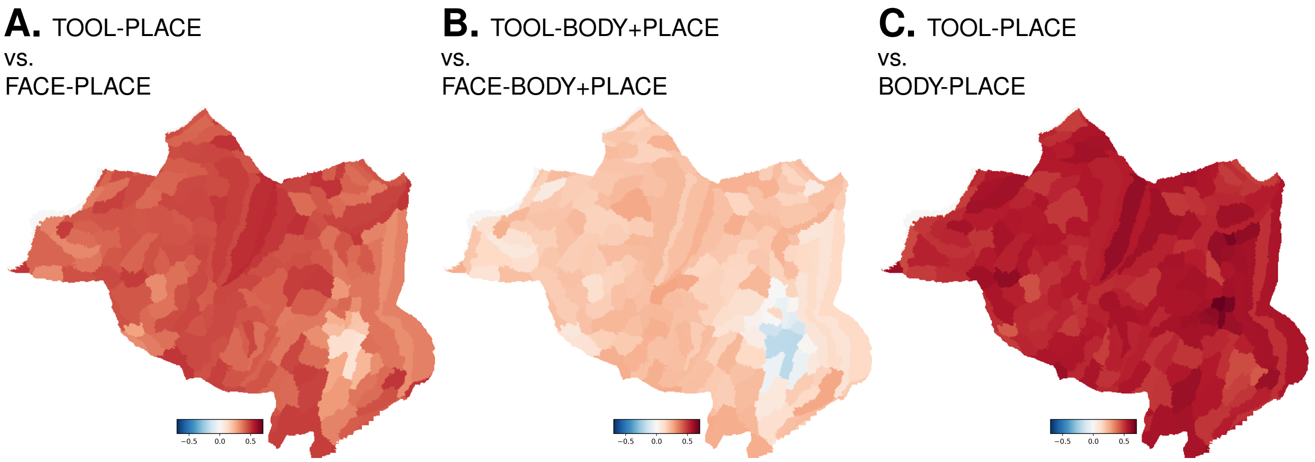


**Figure S8:** Within-experiment parcel-level LI correlations in HCP. This scenario more closely mimics the approach used in the main experiment.

### 6 Long-range face coupling analyses

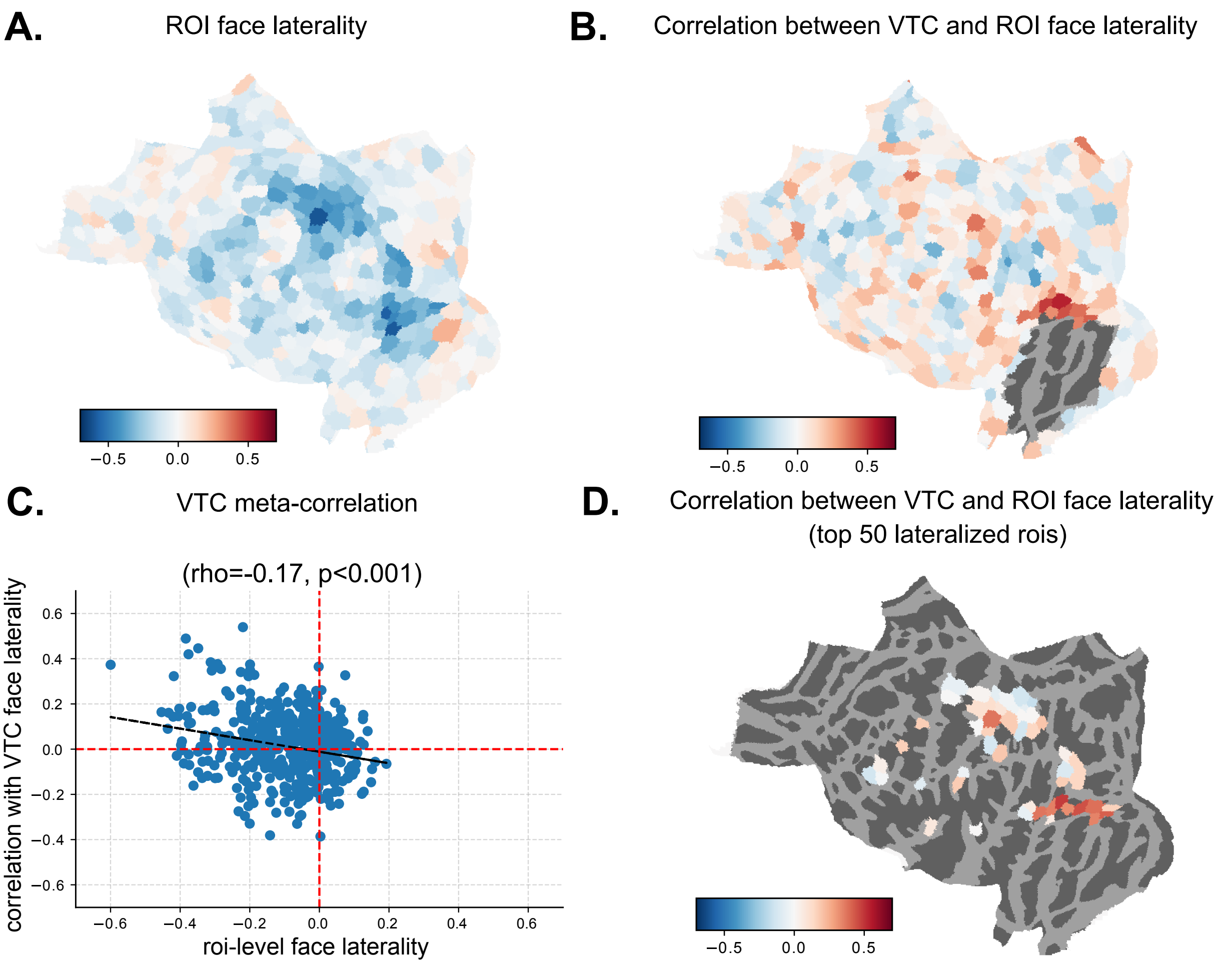


**Figure S9:** Data-driven assessment of whole-brain LI coupling of face selectivity in the main experiment. **A.** Mean of individual atlas parcels across participants. Red corresponds to leftward lateralization, whereas blue corresponds to rightward lateralization. **B.** Correlation of individual patterns of LI in each parcel with that of VTC, across participants, excluding parcels that overlap with the large VTC parcel. **C.** Scatter plot comparing the parcel-level mean LI with the across-participant correlation of LI between the parcel and VTC, along with a line of best fit and result of a spearman correlation across parcels. **D.** Results of **B.** masked to show only the 50 parcels with the largest mean laterality ($|LI|$).

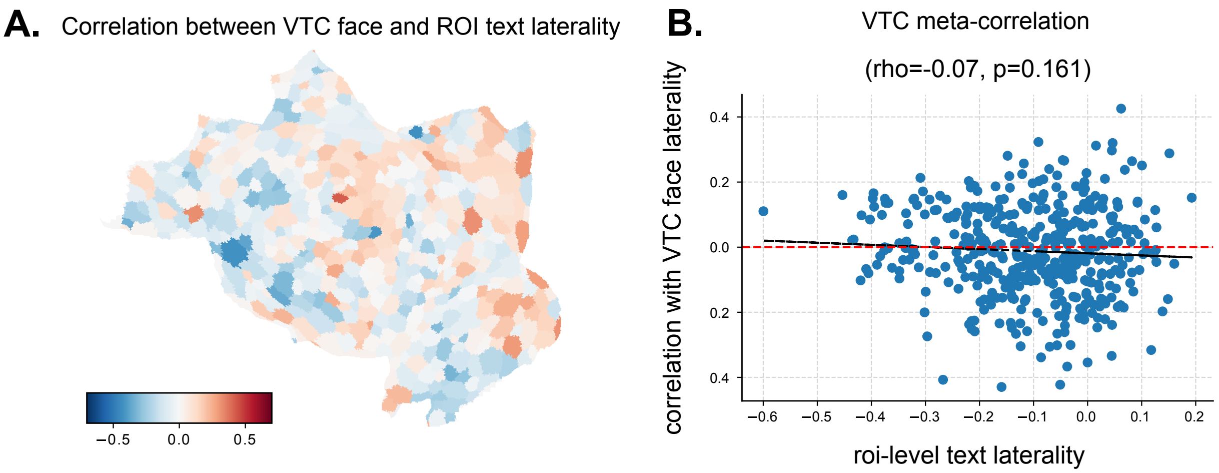


**Figure S10:** Data-driven assessment of whole-brain coupling of text selectivity with VTC face selectivity in the main experiment. **A.** Correlation of individual patterns of text LI in each parcel with face LI in VTC, across participants. **B.** Scatter plot comparing the parcel-level mean LI with the across-participant correlation of LI between the parcel and VTC, along with a line of best fit and result of a spearman correlation across parcels.

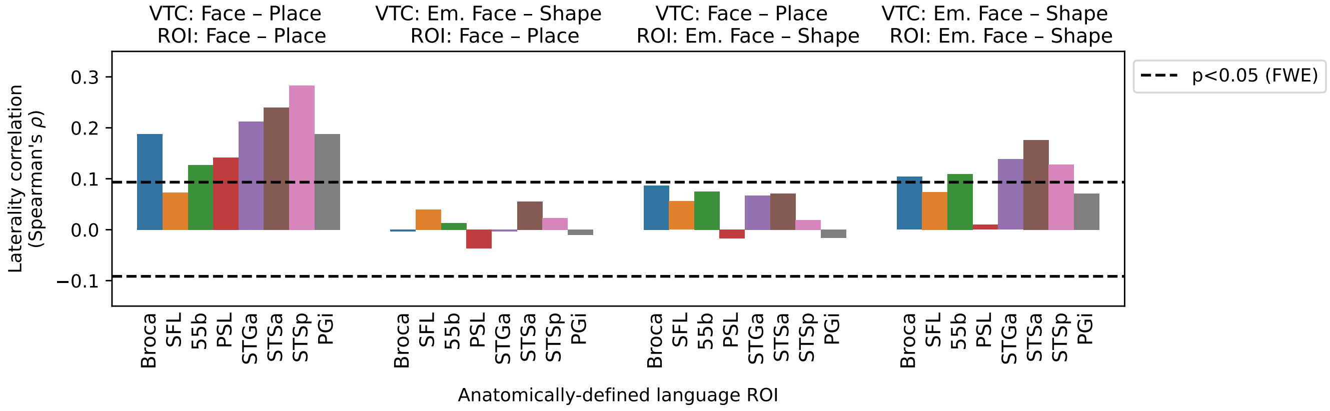


**Figure S11:** LI correlations across face tasks in VTC and Rajimehr language parcels. The same approach as in Figure **??** is used. Laterality coupling with VTC is significant in most parcels in both tasks, however coupling is significantly reduced when comparing across tasks. This suggests that face laterality coupling between VTC and language/social regions may be dependent on the degree of social or emotional processing inherent to the face processing task. One caveat to this result is that we were unable to construct independent subsets of data within each task, so long-range correlations may be partially driven by non-task-based co-fluctuations in the signal.

### References
